# The peculiar photobiology of a giant motile diatom inhabiting the subtidal sediments of the bay of Brest

**DOI:** 10.64898/2026.01.31.703006

**Authors:** Anna Isaia, Gaspard Delebecq, Pauline Breton, Philippe Rosa, Vona Méléder, Aude Leynaert, Johann Lavaud

**Author notes:** E-mail addresses: Pauline Breton, Gaspard Delebecq, Philippe Rosa, Méléder Vona, Aude Leynaert, Johann Lavaud.

## Abstract

Microphytobenthos (MPB) contributes significantly to the marine primary production in estuarine ecosystems. MPB is mainly composed of benthic motile diatoms inhabiting intertidal and shallow subtidal sediments. Unlike intertidal small-sized diatom models, subtidal (≥ 10 m depth) MPB and large-sized (>100 µm) species, have comparatively received much less attention, especially as regards to their photosynthetic productivity. Yet, the subtidal light environment shows unusual (very) low intensities and a green-blue light spectrum at high tide. The present study investigates the light-dependent green-blue responses of the subtidal giant diatom *Pleurosigma strigosum*, combining *in situ* monitoring with laboratory experiments. In both MPB and *P. strigosum*, we documented a strong photophysiological plasticity, and a striking alignment with the photophysiology of (very) low light-adapted polar diatoms. Altogether, our results highlight the nature of subtidal photoadaptation: a very low green-blue light sensitive response, which explains the early blooming of *P. strigosum* at the very beginning of spring and underpins the ecological success of MPB in colonizing coastal subtidal sediments. The general coherence between subtidal *P. strigosum* and MPB light-responses offers a unique model species and growth form to further decipher the specific light-driven metabolism of subtidal MPB under precisely controlled environmental conditions.

## 1 Introduction

Marine coastal ecosystems, particularly estuaries and shallow waters, are home to a wide variety of species and are among the most productive ecosystems, supporting numbers of ecological services (Lebreton et al., 2019; Hope et al., 2020; Serôdio & Paterson, 2022). Their high biological productivity is largely driven by photosynthetic micro-organisms, which supply higher trophic levels and play a central role in biogeochemical cycles (MacIntyre et al., 1996; Lebreton et al., 2019; Hope et al., 2020). While phytoplankton has been extensively studied, the contribution of microphytobenthos (MPB) remains comparatively overlooked, even if it contributes substantially to global carbon cycling, fixing between 30 to 230 g C m^-2^ yr^-1^ on sediment flats (Murray et al., 2019; Bianchi et al., 2024), and MPB productivity exceeding that of phytoplankton in some nearshore waters (Jahnke et al., 2000; Pinckney, 2018). MPB assemblages regroup all microscopic photosynthetic organisms inhabiting the upper intertidal and shallow subtidal sediment layer, and is largely dominated by benthic forms of diatom microalgae (Méléder et al., 2007; Serôdio et al., 2008). The high productivity of these assemblages is remarkable since they thrive in habitats characterized by highly variable and extreme environments, especially in terms of light availability (Kühl & Jorgensen, 1994; Cartaxana et al., 2016; Leynaert et al., 2018).

MPB ecological success under a complex light environment relies on the diatom’s ability to efficiently adjust their photosynthetic productivity to fluctuating and extreme light conditions (Wilhelm et al., 2014; Lepetit et al., 2022). MPB diatoms show two main types of light responses: behavioural (phototaxis) and physiological (photoprotection) mechanisms that allow cells to balance the harvesting and use of light energy, and to prevent and protect from excess light harmful effects (Cartaxana et al., 2011; Serôdio et al., 2012; Barnett et al., 2015; Morelle et al., 2024). Behavioural light response is mediated by directed light-driven cell motility, which allows diatoms to actively move within sediment light gradients (so-called ‘vertical migration’, Serôdio, (2021)), positionate themselves at the light optimum, and avoid prolonged high-light exposure in the sediment upper layer (Consalvey et al., 2004; Serôdio et al., 2023; Jesus et al., 2023). In addition, MPB diatoms employ regulatory processes for physiological photoprotection, especially the non-photochemical quenching (NPQ) of chlorophyll fluorescence that safely dissipates excess light energy (Cartaxana et al., 2011; Barnett et al., 2015; Blommaert et al., 2017). In diatoms, NPQ is controlled by two regulatory components: the LHCx proteins (Buck et al., 2019; Croteau et al., 2025) and the xanthophyll cycle (XC), which involves the light-dependent enzymatic conversion of diadinoxanthin (DD) to diatoxanthin (DT) (Lavaud & Goss, 2014; Giossi et al., 2025). In addition to vertical migration and NPQ/XC, some large-sized (>100 µm length) MPB diatoms show a third type of light response: the complementary light-induced contraction of plastids, named karyostrophy, that was recently revisited in *Pleurosigma strigosum* (Bastos et al., 2025).

MPB research has been strongly biased toward intertidal communities (Hope et al., 2020) and small-sized diatom species (Barnett et al., 2015; Blommaert et al., 2018), including models such as *Seminavis robusta* (Blommaert et al., 2017; Osuna-Cruz et al., 2020; Bondoc-Naumovitz et al., 2025), while shallow subtidal MPB and large-sized benthic diatoms remain comparatively dramatically understudied. This knowledge gap is compounded by (i) the relative scarcity of *in situ* studies, especially for MPB communities found at depths ≥ 10 m (Sundbäck & Jönsson, 1988; Light & Beardall, 2001; Longphuirt et al., 2007; Bourgeois et al., 2010; Chatterjee et al., 2013; Santema & Huettel, 2018), and (ii) the difficulty of isolating and of maintaining large-sized (>100 µm) benthic species in monospecific cultures in lab controlled conditions (De Tommasi et al., 2024; Rizzo et al., 2024; Bastos et al., 2025; Bondoc-Naumovitz et al., 2025).

The bay of Brest (France) offers a great natural laboratory to address this gap. This shallow coastal ecosystem shows a high biomass, diverse (including large-sized genera such as *Pleurosigma*) and productive subtidal MPB community, which biological activity amounts to up to 20% of the total bay primary production (Longphuirt et al., 2007; Chatterjee et al., 2013; Leynaert et al., 2018). These features are remarkable when considering the light environment of subtidal sediments characterised by (very) low surficial irradiance and a light spectrum strongly shifted towards green-blue wavelengths at 10-11 m depth (Chatterjee et al., 2013; Leynaert et al., 2018). In that framework, bay of Brest subtidal MPB was shown, for the first time, to follow strict diel photoperiodic vertical migration in order to optimise its access to light and productivity (Longphuirt et al., 2006). We isolated the large-sized (*ca*. 200 µm) motile diatom *Pleurosigma strigosum* from the subtidal MPB community, one of the first species to grow early during the spring productive season (Leynaert et al., 2018), in order to study its photoadaptive strategy under such a light climate. We found that subtidal *P. strigosum* shows a unique very low green-blue light-adapted response that likely supports its thriving, and the one of MPB diatoms, in subtidal coastal sediments.

## 2 Material and methods

### 2.1 Study site and field monitoring

Lanvéoc (48°17′41 0.23″N – 4°27′12 0.63″W, **Figure 1**), is located in the Southern part of the bay of Brest, a shallow semi-enclosed ecosystem on the Atlantic coast of France. The ecosystem is connected to the Iroise Sea by a narrow strait (2 km wide) and approximately half of its total area lies beneath 5 meters in depth. The Bay has a maximal tidal amplitude of 8 m during spring tides.

**Figure 1:**
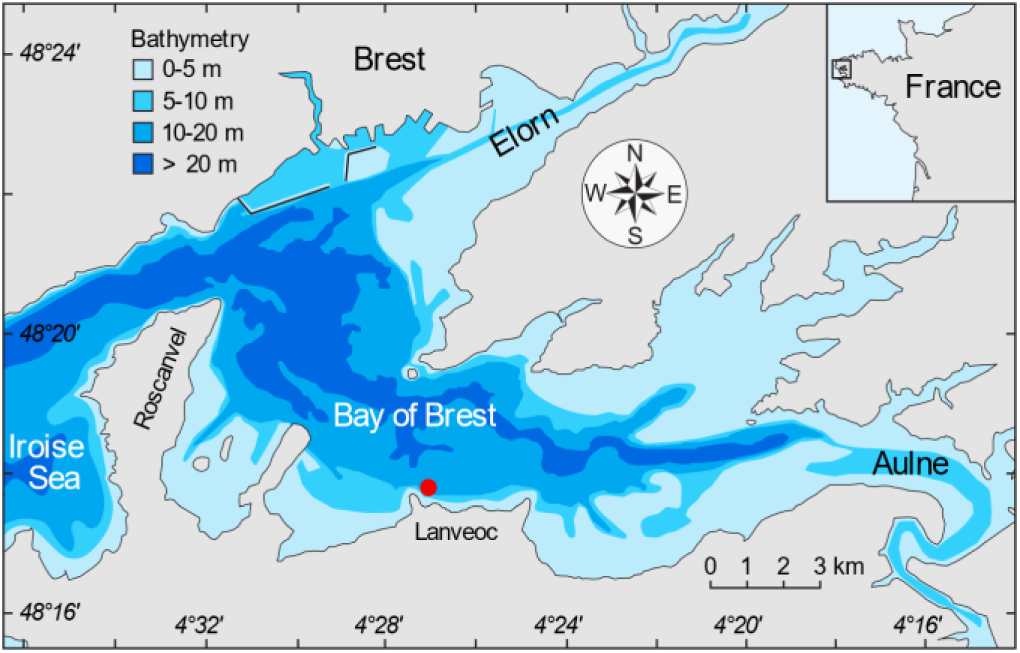
Map of the study site located in Lanvéoc (filled red circle), in the Bay of Brest (modified after Thébault et al., 2022). The inset map (top right) shows the location of the Bay of Brest in France, highlighted by a square. Latitude and longitude grid labels are presented in Degrees-Minutes-Seconds format.

Field monitoring was conducted at high tide biweekly from 2019 to 2025, weather permitting. Sets of physical, chemical and biological measurements were taken at sediment surface or on MPB colonizing Plexiglas plates (10 x 15 cm, Chatterjee et al., (2013), **Figure S1**) placed at the sediment surface. The measurements included: photosynthetic active radiation (PAR, in µmol photons m^-2^ s^-1^), light spectrum, biogenic silica content (BSi, in mmol m^-2^), chlorophyll *a* concentration (Chl *a*, in mg m^-2^), diatom cell abundance (in cells m^-2^) and photophysiological performances. Most parameters were monitored over the period 2019-2024 (PAR, BSi, Chl *a*), with the exception of the light spectra (2019), diatom abundance (2019-23), and photophysiological parameters (2023-25).

PAR and spectrum at the sediment surface were obtained respectively using a PAR sensor equipped on a CTD profiler (SBE 19+ Sea Bird, USA), and a RAMSES (Radiation Measurement Sensor with Enhanced Spectral Resolution, TriOS ACC, Germany) multispectral radiometer with 1 nm resolution between 400-700 nm. Photophysiological parameters were assessed using light curves (LCs) performed directly on the Plexiglas plates with an Imaging-PAM fluorometer (see Section 2.6). For chemical parameters, the diatom biomass was gently scraped off with a standard toothbrush and suspended in a known volume of seawater taken at the sediment surface. Subsamples were taken for Chl *a* and BSi analysis. For Chl *a*, the suspended material was filtered onto GF/F filters (Whatman), and pigments were extracted in 6 mL of 90% acetone. The samples were incubated in the dark at 4 °C for 12 h, centrifuged, and the resulting fluorescence was measured using a Turner Design fluorometer, USA. The equation of Lorenzen, (1966) was used to calculate Chl *a* concentration. For biogenic silica (BSi), samples were filtered onto Nucleopore membrane filters (47 mm diameter) and quantification was based on the method of (Ragueneau et al., 2005) using an auto-analyzer SEAL Analytical® AA3, USA.

### 2.2 Diatom and culture conditions

The diatom *Pleurosigma strigosum* (**Figure S2**) was collected from subtidal sediment sampled in the study site (Lanvéoc) in April 2021, during the MPB bloom (Chatterjee et al., 2013). After isolation, cells were grown in 500 mL Erlenmeyer flasks with a working volume of 200 mL. They were kept in batch culture condition in sterile F/2 enriched seawater (Guillard, (1975), Sigma-Aldrich USA) and additional NaHCO_3_ (80 mg L^-1^ final concentration, Sigma-Aldrich, USA), in a climatic chamber at 15.5 +/- 1°C with a 16:8 hours (light:dark) photoperiod.

### 2.3 Light treatments

Experiments were designed to investigate the photophysiological response of *Pleurosigma strigosum* to its *in situ* light environment, with a focus on light intensity and spectral quality, *i.e.,* spectrum shifted towards green-blue wavelengths (**Figures 2**, **S3**). First, the diatom’s natural light climate was recreated in the laboratory, as close as possible. This was achieved using white LEDs (LXML-PD01, 4100 K; LUXEON REBEL, LUMILEDS, Aachen, Germany) covered with a green filter (LEE, Fern Green FL 122) to recreate the green spectral contribution, alongside with blue LEDs (440 nm, SL3500-E Photons Systems instruments, Czech Republic) to provide the necessary blue light contribution. A gradient of six increasing green-blue (GB) light intensities was selected from 2 to 170 µmol photons m^-2^ s^-1^ encompassing average *in situ* light intensities measured along the year at the study site (see **Figure 3**). Light experiments were performed after a pre-photoacclimatation period of at least 14 days under the corresponding light regime. Samples for analyses and measurements were harvested from cultures in their exponential growth phase.

**Figure 2:**
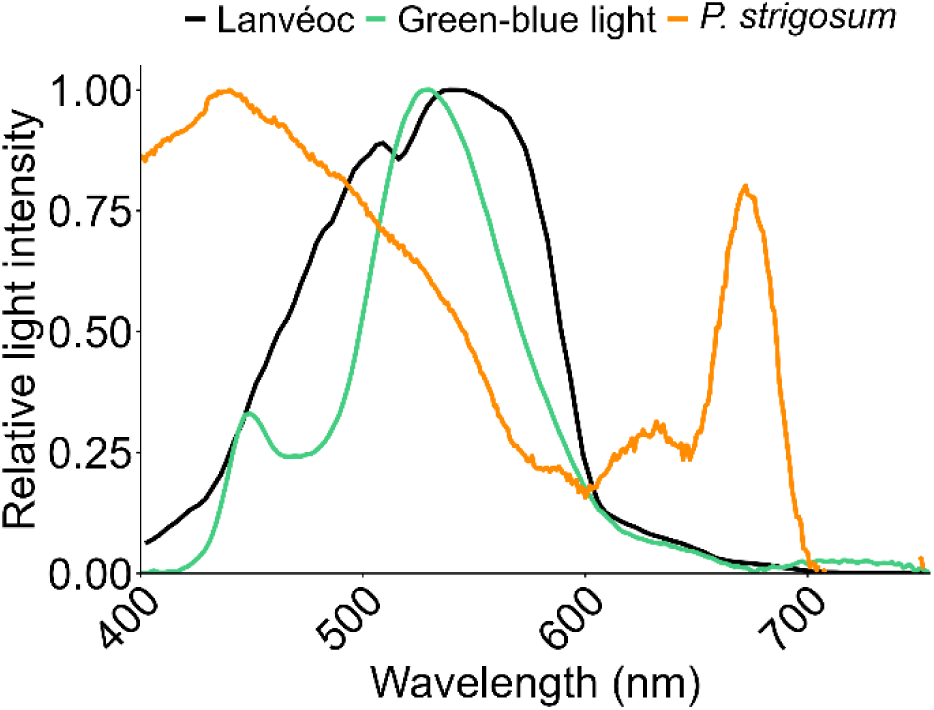
Light spectrum at the study site (Lanvéoc, corresponds to the mean trace in **Figure S3**), emission spectra of the green-blue experimental light source and the typical absorption spectrum of *Pleurosigma strigosum* cells.

**Figure 3:**
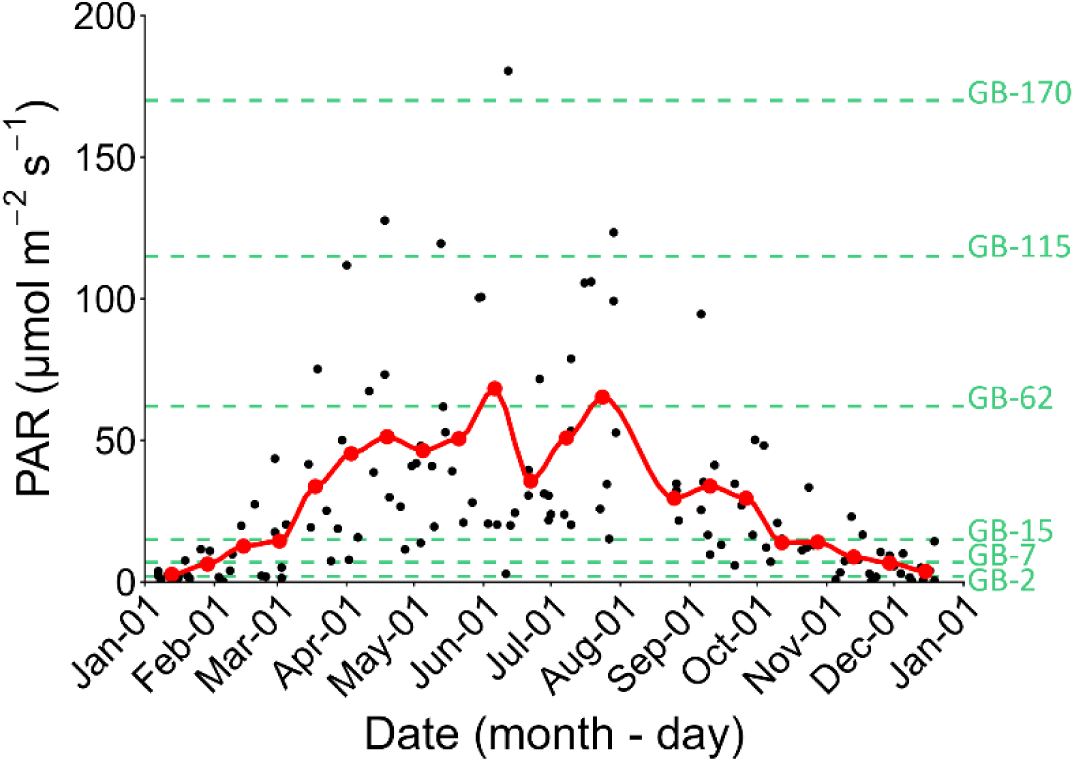
Annual profile of the Photosynthetically Active Radiation (PAR) at the sediment surface of the study site (11.9 ± 1.4 m depth); the black dots show raw data, the red dots and line represent the two weeks mean; green dashed lines show the six growth light conditions used in this study: 2, 7, 15, 62, 115 and 170 µmol photons m^-2^ _s-1._

### 2.4 Growth performances

Cell growth was followed by cell counting using a gridded Sedgewick Rafter counting chamber under a binocular magnifier. Specific growth rates (µ, day^-1^, see **Table S1** for the definition and units of all parameters used in this study) were calculated during the logarithmic growth phase using the following equation:

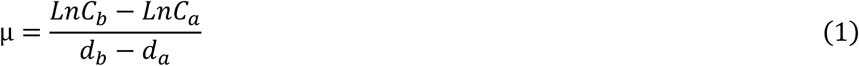

where 𝐶_𝑎_ and 𝐶_𝑏_ represent cell concentrations (cells mL^-1^) at days 𝑑_𝑎_ and 𝑑_𝑏_. Growth rate parameters were estimated following the approach of Croteau et al., (2022), by fitting the model developed by Eilers & Peeters, (1988) to the experimental data, using the equation:

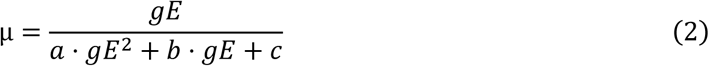

where, growth rate parameters were estimated using the following equations:

-the maximum light use efficiency for growth, α_µ_ in d^-1^ µmol photons m^-2^ s^-1^:

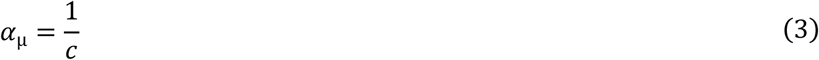

-the maximal growth rate, µ_max_ in day^-1^:

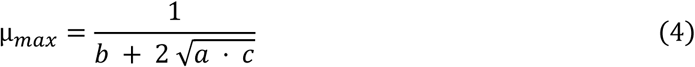

-the light intensity for growth rate saturation, K_E_ in µmol photons m^-2^ s^-1^:

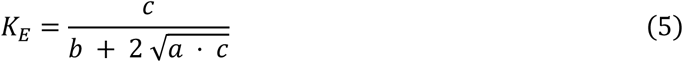

-the optimal growth intensity, gE_µmax_ in µmol photons m^-2^ s^-1^:

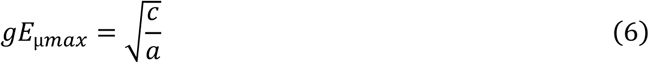

The frustule cell size was measured using a scanning electron microscope (SEM, JCM-7000 NeoScope, Switzerland). Cells were treated with hydrogen peroxide (H₂O₂) to remove organic matter, and subsequently dehydrated through a graded ethanol series (15%, 30%, 50%, 70%, 90%, 95%, and 100%; 10 min at each concentration). Frustules were then collected on polycarbonate filters (25 mm diameter, 0.2 µm pore size, Cytiva, USA) and sputter-coated with a conductive metal layer prior to SEM observation. For each sample, at least 10 cells were measured for both length and width (µm).

### 2.5 Elemental analyses

Prior to sample collection, all equipment was pre-combusted in a furnace (4 h, 450 °C) to prevent carbon and nitrogen contamination. Carbon and nitrogen content (QC and QN) were measured using 10 mL of culture filtered through glass fibre filters (Whatman GF/F 25 mm, 1.2 µm pore size, GE HealthCare, China). Filters were dried at 60°C for at least 24 h prior to analysis.

Elemental composition was measured using an elemental analyser (Thermo Fischer Flash EA 1112, USA), following the combustion method of (Strickland & Parsons, 1972). The C:N ratio was calculated as:

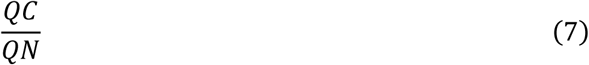

The daily carbon (C) production (in ng cell^-1^ day^-1^) was calculated as:

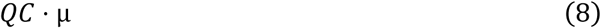

To determine the BSi, 60 cleaned cells (*i.e.,* cleaned from culture medium with artificial seawater) were placed onto a polycarbonate membrane (Whatman Nucleopore 47 mm, 0.6 μm pore size, Cytiva, USA). The quantification of BSi was as in Study site and field monitoring.

### 2.6 Photosynthetic performances

To assess the photophysiological state of *in situ* (Lanvéoc) MPB (see Section 2.1) and of *P*. *strigosum* cultures, the IMAGING-PAM (PAM: pulse amplitude modulated) fluorometer was used. The IMAGING-PAM presents the dual advantage of performing, with the same light curve (LC) protocol, measurements on the MPB colonised Plexiglas plates (see Section 2.1) and on *P. strigosum* cells deposited at the bottom of 12 wheel-plates, which is the closest to their *in situ* arrangement in biofilm at the surface of sediment. The IMAGING-PAM fluorometer (Maxi-PAM M-series IMAG-MAX/L ‘blue version’, 450 nm, see Barnett et al., (2020)) was used to measure Chl *a* fluorescence for an overall determination of photophysiological parameters. Measurements were performed on samples dark-acclimated for at least 30 min. LCs were generated using 20 incremental light intensities of 60 s, ranging from 6.4 to 867 µmol photons m^-2^ s^-1^. The maximum quantum efficiency of PSII, Fv/Fm, was estimated according to the equation (Schreiber et al., 1995):

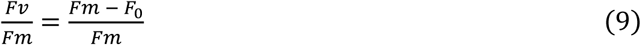

with 𝐹𝑣 the variable fluorescence, 𝐹𝑚 the maximum fluorescence and 𝐹_0_ the minimum fluorescence yield (dark-acclimated minimum fluorescence yield). LCs were used to build relative photosynthetic electron transport rate (rETR) *versus* light intensity (PAR) curves. They were fitted with the (Eilers & Peeters, 1988) model to estimate the maximum relative photosynthetic electron transport rate (rETRm), the optimal light intensity for reaching rETRm (Eopt, in µmol photons m^-2^ s^-1^) and the light intensity for saturation (Ek, in µmol photons m^-2^ s^-1^).

In parallel, the regulated non-photochemical quenching yield (Y(NPQ)) was calculated to evaluate photoprotective heat dissipation. It was computed according to (Klughammer & Schreiber, 2008):

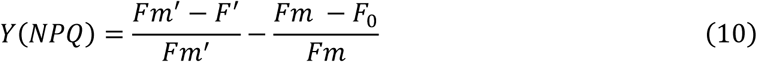

where 𝐹𝑚’ is the maximum fluorescence in the light-adapted state, 𝐹𝑚 is the maximum fluorescence in the dark-acclimated state, 𝐹’ the fluorescence yield in the light, 𝐹_0_ the minimum fluorescence yield (dark-acclimated minimum fluorescence yield). To further quantify photoprotective capacity, the model developed by (Serôdio & Lavaud, 2011) was applied to extract key parameters: the maximum Y(NPQ), Y(NPQm), and the intensity for half-saturation of Y(NPQm) (Y(E50_NPQ_), in µmol photons m^-2^ s^-1^).

### 2.7 Pigment analysis

Pigment composition was estimated by High Performance Liquid Chromatography (HPLC) as in Méléder et al., (2003). Briefly, aliquots of *P. strigosum* cultures (10 to 20 mL) were filtered on Isopore membrane filters (1.2 µm, Merck Millipore, Germany), directly frozen in liquid nitrogen and stored at −80°C until analysis. Pigments were extracted using 2 mL of methanol at 95% (MeOH/H2O v/v) with 2% of ammonium acetate, and an internal standard, the trans-β-Apo-8’-carotenal at 1 mg L^-1^ final concentration (usp, USA). Each sample was mixed, stored at −20 °C for 15 min and syringe filtered (0.2 µm, Whatman, Avantor, USA). Extracts were analysed as described by Mantoura & Llewellyn, (1983) and adapted by Méléder et al., (2003). Quantification (in pg cell^-1^) was carried out at 440 nm by comparison with pigments standards (DHI, Denmark).

### 2.8 Statistical analyses

Statistical analyses were performed with R studio version 4.3.3 (2024-02-29). Data distribution was assessed using the Shapiro-Wilk test, and homogeneity of variances was evaluated with the Levene’s test. Based on these results, either a parametric one-way ANOVA or a non-parametric Kruskal-Wallis test and *post hoc* parametric Student’s *t*-test or a non-parametric Wilcoxon rank-sum test was applied to compare light treatments. The statistical significance was determined at 95% of confidence level (*p value* < 0.05). linear and non-linear regression were fitted to raw data with *geom_smooth* function in *ggplot2*. In addition, *stat_regline_equation* (package *ggpmisc*) was used to produce equations and R². For linear regression, correlation test was applied using *cor.test* function.

## 3 Results

### 3.1 Lanvéoc subtidal sediments environmental context

#### 3.1.1 In situ light conditions

Photosynthetically active radiation (PAR) at the sediment surface (11.9 ± 1.4 m depth) followed a typical seasonal pattern (**Figure 3**, black dots). From January to early March, PAR values were below 10 µmol photons m^-2^ s^-1^ with a minimum of 0.19 µmol photons m^-2^ s^-1^. From March to June the PAR increased sharply, reaching a maximum of 180 µmol photons m^-2^ s^-1^, after which it progressively declined below 10 µmol photons m^-2^ s^-1^ from October to December. Based on the biweekly average PAR (red line **Figure 3**), we determined the reference intensities that were further used to set-up our experimental green-blue light treatment (Section 3.2), including minimum and maximum intensities (2 and 170 µmol photons m^-2^ s^-1^, respectively) and four intermediate intensities encompassing all seasons (see the dashed lines, **Figure 3**). The year-round sediment surface spectrum was broadly constant, showing maximal intensity in the green-blue 490-560 nm range (**Figure S3**), and the mean year-round spectrum (dashed line, **Figure S3**) was further used to set-up our experimental green-blue light treatment (Section 3.2 and **Figure 2**).

#### 3.1.2 In situ microphytobentos (MPB) abundance

Total Chl *a* concentrations were very variable (6.14 ± 4.99 mg m^-2^) with a maximum of 34.69 mg m^-2^ early April and secondary weaker peaks late May (29.35 mg m^-2^) and late October (15.43 mg m^-2^) (**Figure 4**). Regarding the MPB in itself, diatom cells abundance was also very variable with a maximum of 1.96 × 10⁹ cells m^-2^ in early April and secondary peaks of abundance (*ca*. 1 × 10⁹ cells m^-2^) in mid-April, early May, and late-June/mid-July. The MPB community was dominated by three genera, namely *Navicula*, *Fragilaria* and *Amphora* (**Figure S4**). The *Pleurosigma* genus was present throughout the year with an average abundance of 7.88 ± 4.63 × 10^4^ cells m^-2^ year^-1^ (<1% of the total abundance). The biweekly average BSi concentration showed an average value of 4.34 ± 1.66 mmol m^-2^ with a maximum of 7.91 mmol m^-2^ early April and secondary maxima in late June-July and in September-October.

**Figure 4:**
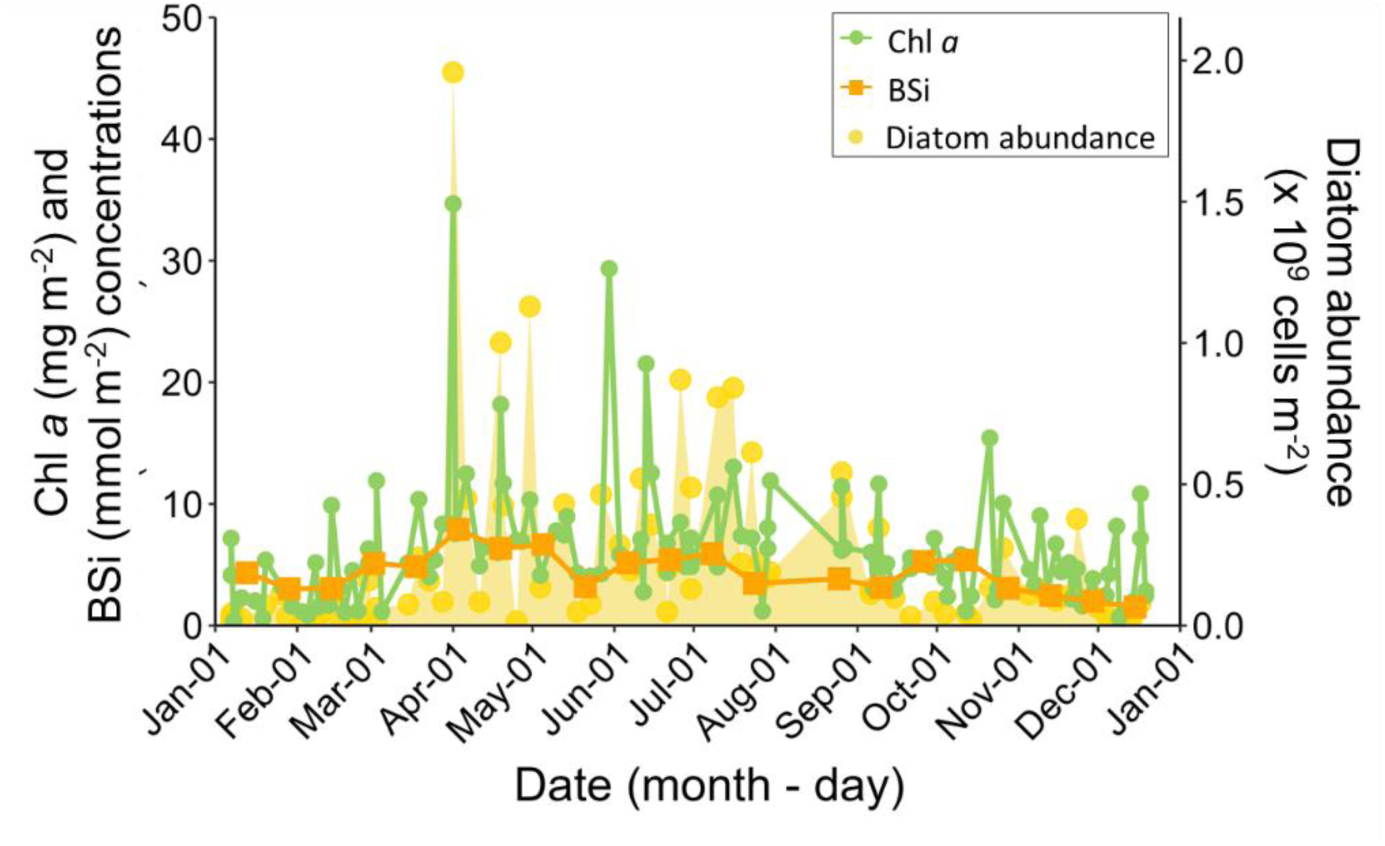
Annual profile of chlorophyll *a* (Chl *a*, green dots and line), biogenic silica (BSi, orange squares and line), and diatom abundance (yellow dots and area) of microphytobenthos at the study site. Chl *a* and diatom abundance are shown as raw data and BSi as the two weeks mean. The main taxonomic information behind total diatom abundance is found in **Figure S4**.

#### 3.1.3 In situ MPB photophysiology

The MPB maximum quantum efficiency of PSII, Fv/Fm (see **Table S1** for definition of all photophysiological parameters), which is a robust index of the ‘health’ status of cells, was fairly high for MPB and steady throughout the year (0.55 ± 0.09). In contrast, the maximum relative photosynthetic electron transport rate (rETRm), *i.e.,* the potential of MPB for photochemistry, and the associated light parameters (Ek, the light intensity for saturation of photochemistry, and Eopt, the optimal light intensity for photochemistry, *i.e*., for reaching rETRm) exhibited strong seasonal dynamics, with minima in winter (with values as low as 1.61 ± 0.38) and 15 times higher maxima from the end of May to mid-July (**Figure 5a**; **Figure S5a, c**). Noteworthy, Ek and Eopt maxima (Ek: 50-75 µmol photons m^-2^ s^-1^; Eopt: 150-220 µmol photons m^-2^ s^-1^) well matched with the range of *in situ* irradiances (**Figure 3**), including the biweekly average PAR maximum (65-70 µmol photons m^-2^ s^-1^, red line **Figure 3**) and the recorded maximum PAR (180 µmol photons m^-2^ s^-1^, black dots **Figure 3**). The direct relationships between rETRm, Ek, Eopt and seasonal PAR followed a second-degree polynomial regression (**Figures 5b, S5b, d**) and allowed the estimation of the irradiance corresponding to their maxima (PARmax), which were close: respectively 122, 114 and 113 µmol photons m^-2^ s^-1^ for rETRm, Ek and Eopt respectively (**Table S2**). Also noteworthy, was the relative steadiness of the ratio Eopt/Ek throughout the year with no apparent seasonal pattern (2.53 ± 0.52). The maximal regulated non-photochemical quenching yield, Y(NPQm), which illustrates the potential of MPB for photoprotection was also mainly stable (*ca*. 0.59) throughout the year and across the PAR range (**Figure 5c, d**), albeit reaching a maximum value of 0.92 ± 0.06 in April and its lowest values (*ca*. 0.45-0.50) during the winter months (**Figure 5c**). Because the light intensity for half-saturation of Y(NPQm) (*i.e*., for reaching 50% of Y(NPQm), Y(E50_NPQ_)) was also steady throughout the year and across PAR range (*ca*. 426 µmol photons m^-2^ s^-1^), the ratio Y(E50_NPQ_)/Eopt, which illustrates the relationship between photoprotection and photochemistry, fitted to an inverse seasonal pattern, with the highest values in winter (*ca*. 20-25) and the lowest, closest to 1, values in spring/summer (*ca*. 1.5-2) (**Figure 5e**). This trend was also well captured by its relationship with PAR (power regression **Figure 5f**), with a sharp decrease at low PAR values (almost divided by two from PAR 1 to 10 µmol photons m^-2^ s^-1^) followed by a more gradual decline at higher intensities, reaching a minimum at PAR 100 µmol photons m^-2^ s^-1^ (**Table S2**), strikingly close to the PARmax of rETRm, Ek and Eopt. Also, noteworthy, all fits (**Figure 5a, e**; **Figure S5a, c**) proposed very close dates for reaching maximal (rETRm-June 28, Ek-June 25, Eopt-June 23) and minimal (Y(E50_NPQ_)/Eopt-June 20) values.

**Figure 5:**
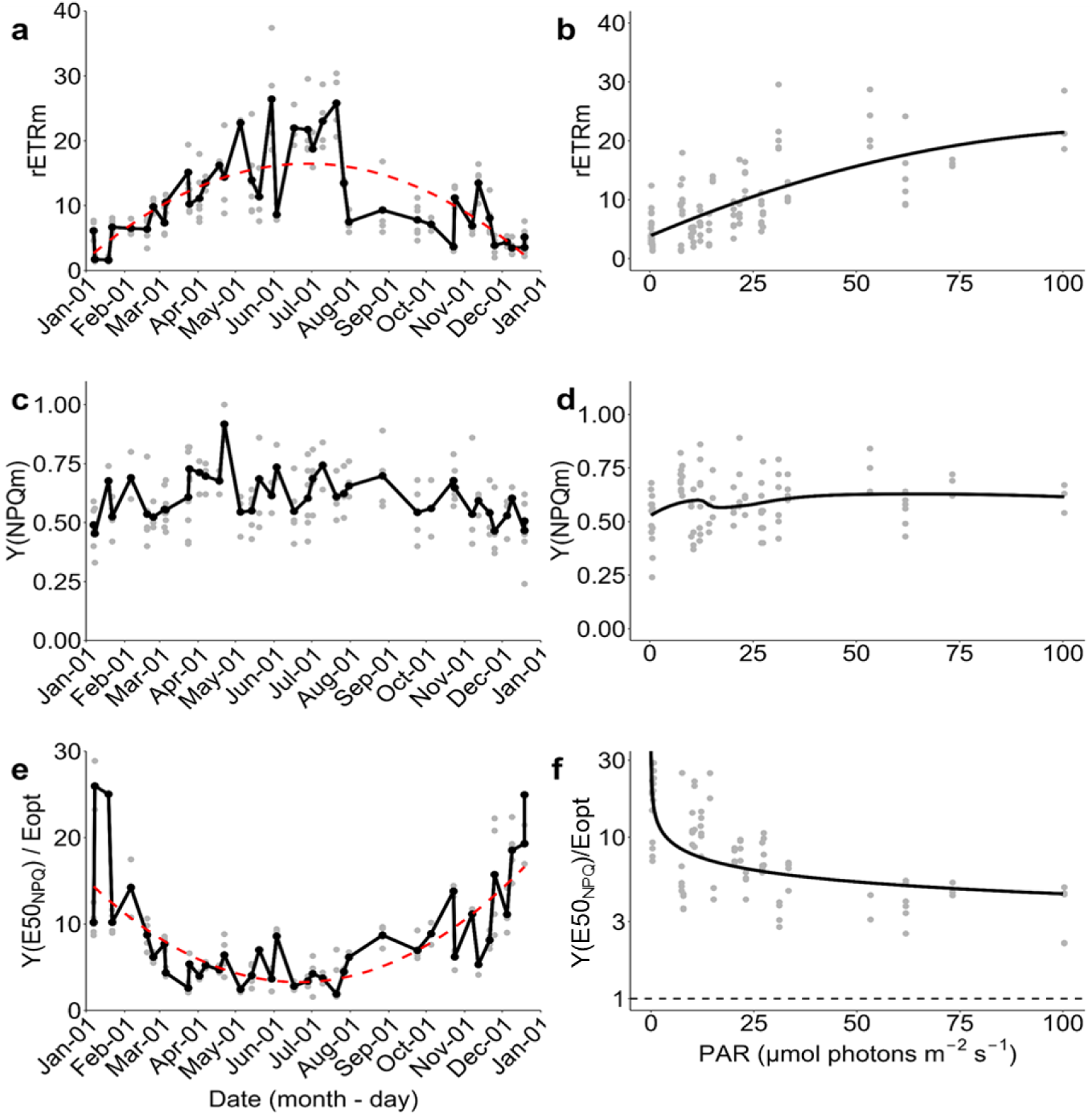
Annual dynamics of microphytobenthos (MPB) photophysiological parameters at the study site and their relationship with the corresponding photosynthetically active radiation (PAR) at the sediment surface. (**a**) Annual maximum relative photosynthetic electron transport rate (rETRm); (**b**) rETRm *versus* the corresponding PAR; (**c**) annual quantum yield of maximal regulated non-photochemical quenching Y(NPQm); (**d**) Y(NPQm) *versus* corresponding PAR; (**e**) annual ratio between the light intensity for which Y(NPQ) reaches half-maximal saturation and Eopt, the light intensity for reaching rETRm (Y(E50_NPQ_)/Eopt) and (**f**) Y(E50_NPQ_)/Eopt *versus* corresponding PAR. Grey dots indicate raw data. In (**a**), (**c**) and (**e**), black dots and solid lines represent the mean of daily measurements, and red dashed lines, in (**a**) and (**e**) indicate second-order polynomial regressions (R² = 0.49 and 0.55 for (**a**) and (**e**) respectively). In (**b**) and (**f**), black curves represent regression and smoothing models fitted to raw data (second-order polynomial regression for rETRm, R² = 0.49; power regression for Y(E50_NPQ_)/Eopt, R² = 0.44) and dashed line in (**f**) is 1:1 ratio (Y(E50_NPQ_)/Eopt).

The ratios of light acclimation and photoprotection parameters to *in situ* irradiance were examined to explore the MPB photoadaptation to its light regime (**Figure 6**). All three ratios followed a logarithmic trend declining with increasing PAR. The ratio Ek/PAR dropped below 1 at low irradiance (*ca.* 26 µmol photons m^-2^ s^-1^ according to the regression **Figure 6a**), indicating that the light saturation point of MPB was reached well before the maximum available PAR in the field. In contrast, the ratio Eopt/PAR decreased more gradually, reaching 1 at higher PAR value *ca.* 96 µmol photons m^-2^ s^-1^ (**Figure 6b**). Interestingly, the ratio Y(E50_NPQ_)/PAR remained consistently above 1 across the irradiance range, although it declined progressively with increasing light (**Figure 6c**).

**Figure 6:**
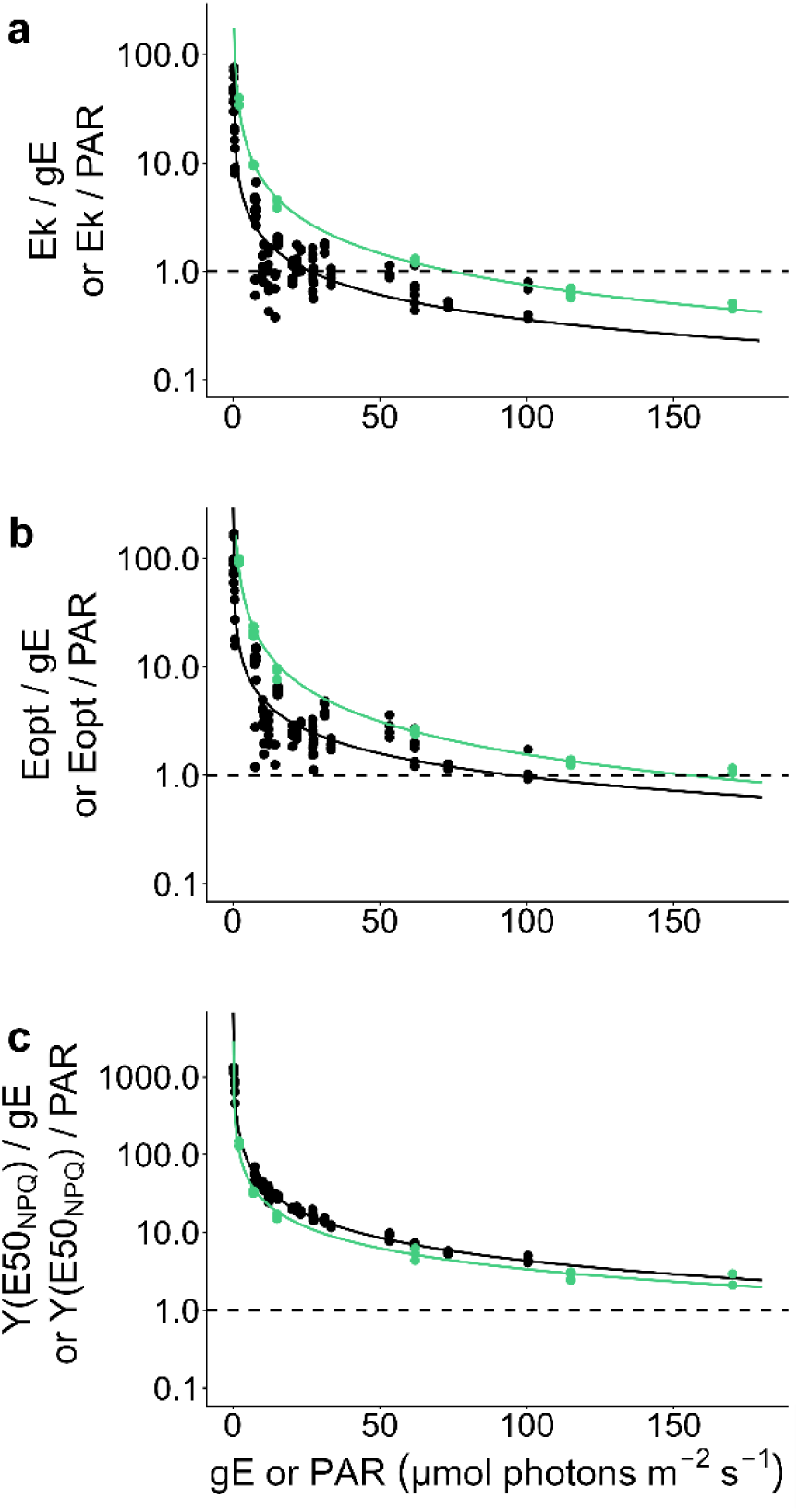
Ratios of light use optima (see **Table S1** for definitions) and the corresponding PAR (photosynthetically active radiation) at the sediment surface of the study site for MPB (black dots) or the corresponding growth light intensity (gE) for *P. strigosum* (green dots) over PAR and gE respectively. (**a**) Ek/PAR and Ek/gE, (**b**), Eopt/PAR and Eopt/gE, and (**c**) Y(E50_NPQ_)/PAR and Y(E50_NPQ_)/gE. Dots are raw data, solid lines are power regression models fitted to the raw data, R² for MPB are (**a**) 0.84, (**b**) 0.83 and (**c**) 0.99 and for *P. strigosum* (**a**) 1, (**b**) 0.99 and (**c**) 0.98. Dashed lines are 1:1 ratios.

### 3.2 Growth and photoacclimation of *Pleurosigma strigosum* to a green-blue light spectrum

In order to understand how *P. strigosum* responds to its specific *in situ* light environment, we grew it under an experimental light spectrum recreating at best the mean year-round spectrum at the surface of sediments (**Figure 2** based on **Figure S3**), and encompassing six growth light intensities (gE) representative of the PAR seasonal variations (**Figure 3**).

#### 3.2.1 Growth performances and elemental analysis

The specific growth rate (µ) of *P. strigosum* followed a light intensity-dependent saturating curve (R² = 1, **Figure 7a**) with a light intensity for growth rate saturation, K_E_, of 47 µmol photons m^-2^ s^-1^. The maximal growth rate (µ_max_, 0.37 day^-1^) was reached for a gE of 117 µmol photons m^-2^ s^-1^ (gE_µmax_, see **Table S2**). In parallel, Fv/Fm was stable over the light gradient (0.60 ± 0.042, **Figure 5a**, **Table S3**). Frustule morphometrics showed a mean cell length of 198 ± 10 µm and a mean width of 24 ± 0.7 µm with significantly slightly longer cells (*ca.* 205 µm) for 15 and for 62 µmol photons m^-2^ s^-1^ (**Tables S3, S4**).

**Figure 7:**
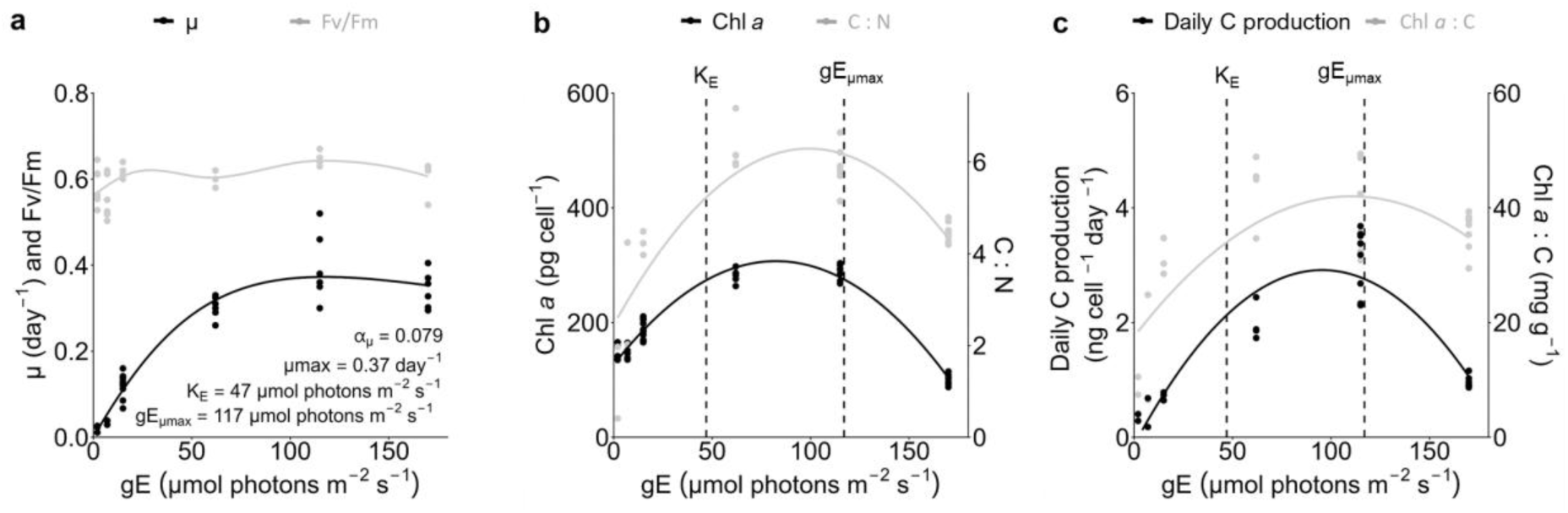
(**a**) Specific growth rates, µ, and the maximum quantum efficiency of PSII, Fv/Fm, (**b**) Cellular chlorophyll *a* (Chl *a*) and Carbon to Nitrogen ratio (C : N), (**c**) chlorophyll *a* to C ratio (Chl *a* : C) and daily carbon (C) production of *P. strigosum* cells grown under increasing green-blue growth light intensities (gE). Dots are raw data. Lines represent model fits or smoothing applied to raw data: (**a**) Eilers & Peeters, (1988) model (R² = 1), (**b**) second-order polynomial regression for Chl *a* (R² = 0.95) and for C : N (R² = 0.75), (**c**) second-order polynomial regression for daily C production (R² = 0.83), and for the Chl *a* : C ratio (R² = 0.52). Dashed lines in (**b**) and (**c**) indicate gE = K_E_ (47 µmol photons m^-2^ s^-1^, the light saturation intensity for growth) and gE = gE_µmax_ (117 µmol photons m^-2^ s^-1^, the light at which the growth rate is maximal).

While the C cell content showed a logarithmic decrease with gE (**Figure S6**), the Chl *a* cell content, carbon-to-nitrogen (C:N) ratio, Chl *a*-to-C ratio (Chl *a*:C), that illustrates the cells capacity to fix C *versus* their light harvesting capacity, and the daily C production followed a similar hyperbolic trend (**Figure 7b, c**). The fitting second-order polynomial models (**Figure 7b, c**) provided close gEmax for reaching maxima: 82, 99, 110 and 96 µmol photons m^-2^ s^-1^ for Chl *a*, C:N, Chl *a*:C and C production, respectively (**Table S2**). When these maxima are normalized to gEµmax, the closest to 1 was Chl *a*:C (0.94) while the Chl *a* content maximum was already reached for 70% of gE_µmax_ (**Table S2**). Noteworthy is the fact that the mathematical fittings (**Figure 7**) extrapolated a high Chl *a* (126 pg cell^-1^) and C (0.63 pg C µm^-3^) cell contents at 0 µmol photons m^-2^ s^-1^, for an expected null growth rate and a negative daily C production (see **Table S5**). Nevertheless, even for gE as low as 2 µmol photons m^-2^ s^-1^, *P. strigosum* was able to generate a relevant daily C production of 0.34 ± 0.08 ng C cell^-1^ d^-1^ (**Figure 5c**).

#### 3.2.2 Accessory pigment content

Accessory cell pigment contents also followed second-order polynomial functions (**Figure 8**) with an average fitted gEmax of 82 ± 3 µmol photons m^-2^ s^-1^, a value similar to the Chl *a* cell content, and very close to the C:N ratio and the daily C production (**Table S2**). When expressed relative to Chl *a*, pigment ratios were broadly stable across gE (**Figure S7**, **Table S6**). Similarly, to Chl *a*, the cell pigment content of accessory pigments was predicted to be high at 0 µmol photons m^-2^ s^-1^, especially fucoxanthin (78 pg cell^-1^) (**Table S5**).

**Figure 8:**
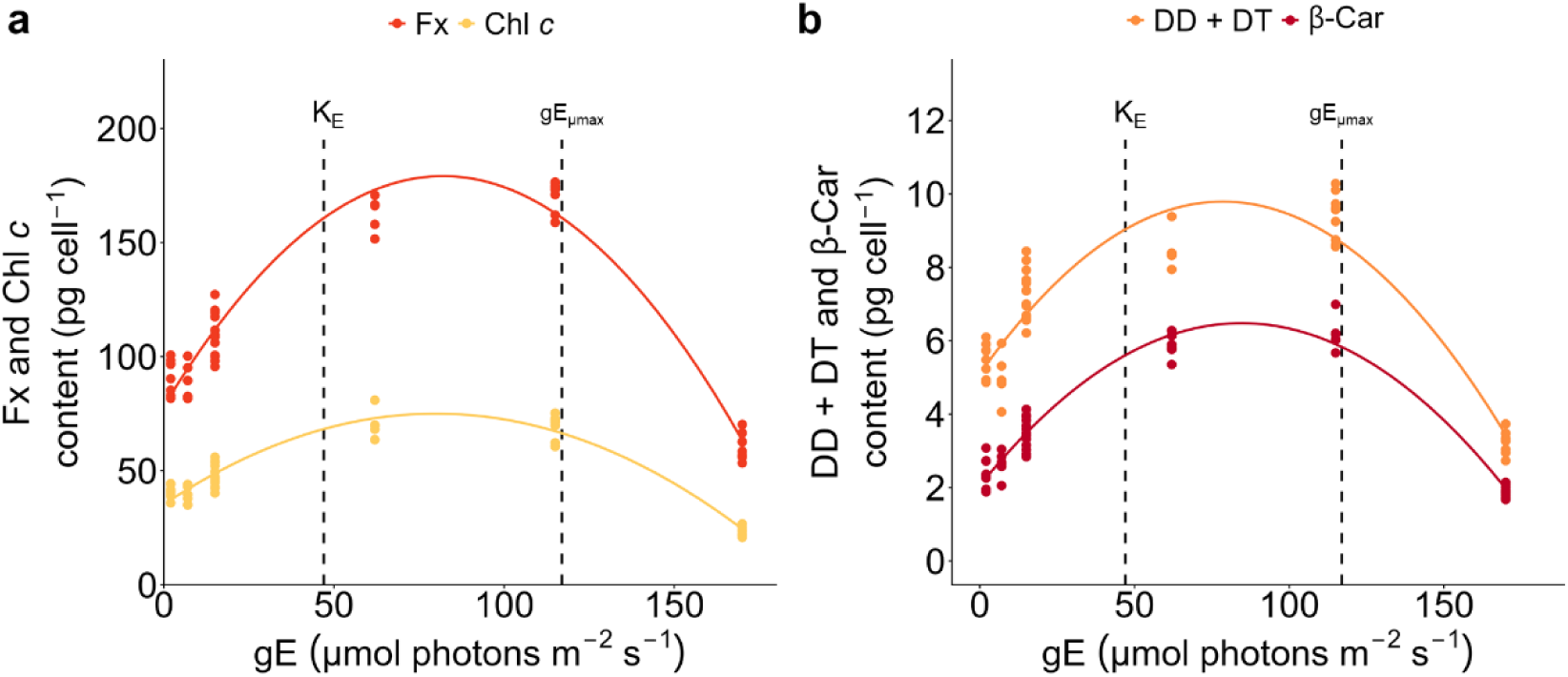
Pigment content of *P. strigosum* cells grown under increasing green-blue growth light intensities (gE). Pigments are (**a**) fucoxanthin (Fx), chlorophyll *c* (Chl *c*), and (**b**) the xanthophyll cycle pigments (DD+DT, diadinoxanthin and diatoxanthin), and β-carotene (β-Car). Dots are raw data. The curves are fitted using second-order (polynomial) regression model; corresponding R² are 0.94, 0.92, 0.85 and 0.95 for Fx, Chl c, DD+DT and β-Car, respectively. Dashed lines indicate gE = K_E_ (47 µmol photons m^-2^ s^-1^, the light saturation intensity for growth) and gE = gE_µmax_ (117 µmol photons m^-2^ s^-1^, the light at which the growth rate is maximal). The corresponding data expressed in mol 100 mol Chl a^-1^ are found in **Figure S8**.

#### 3.2.3 Photophysiology as a function of light intensity

In order to decipher the photophysiological response of *P. strigosum*, we investigated the same main parameters with the same PAM fluorometer and light curve protocol as for the *in situ* MPB: rETRm, Y(NPQm), the ratio Y(E50_NPQ_)/Eopt, and their corresponding light intensities optima (see Section 3.1). rETRm showed values systematically higher than those of *in situ* MPB (*ca.* 40 *versus* 25-27; **Figure 9a** and **Figure 5a, b**) and a fitted gEmax of 110 µmol photons m^-2^ s^-1^ close to the one of µmax (*i.e*., gEmax_rETRm_/gE_µmax_ = 0.94) and daily C production, and to the ones of MPB rETRm (**Table S2**). Light intensities for photosynthetic performances, Ek (72 ± 7 µmol photons m^-2^ s^-1^) and Eopt (154 ± 19 µmol photons m^-2^ s^-1^) were in the upper range of *in situ* MPB values (see **Figure S5**) and were contrastingly stable over gE. Nevertheless, the Eopt/Ek ratio was very similar to the one of *in situ* MPB (2.25 ± 0.28 *versus* 2.53 ± 0.52 for *in situ* MPB). Y(NPQm) values were in the exact same range as *in situ* MPB values, *i.e.,* from 0.50 to 0.90 (**Figure 9b** and **Figure 5c, d**) increasing with gE through a power law relationship (**Figure 9b**). Fitted gEmax was estimated at 204 µmol photons m^-2^ s^-1^, *ca.* two times higher than gEmax_rETRm_ (**Table S2**). Different from *in situ* MPB, Y(E50_NPQ_) linearly increased with gE (**Figure S8**) and with values similar to *in situ* MPB only for the highest gE (433 ± 66 at 170 µmol photons m^-2^ s^-1^ *versus* an average of 431 ± 53 µmol photons m^-2^ s^-1^ for MPB). The Y(E50_NPQ_)/Eopt ratio was in the range of the lowest values for *in situ* MPB (1.39 to 2.30, see **Figure 9c** and **Figure 5e, f**) and increased over gE through a power law relationship (**Figure 9c**).

**Figure 9:**
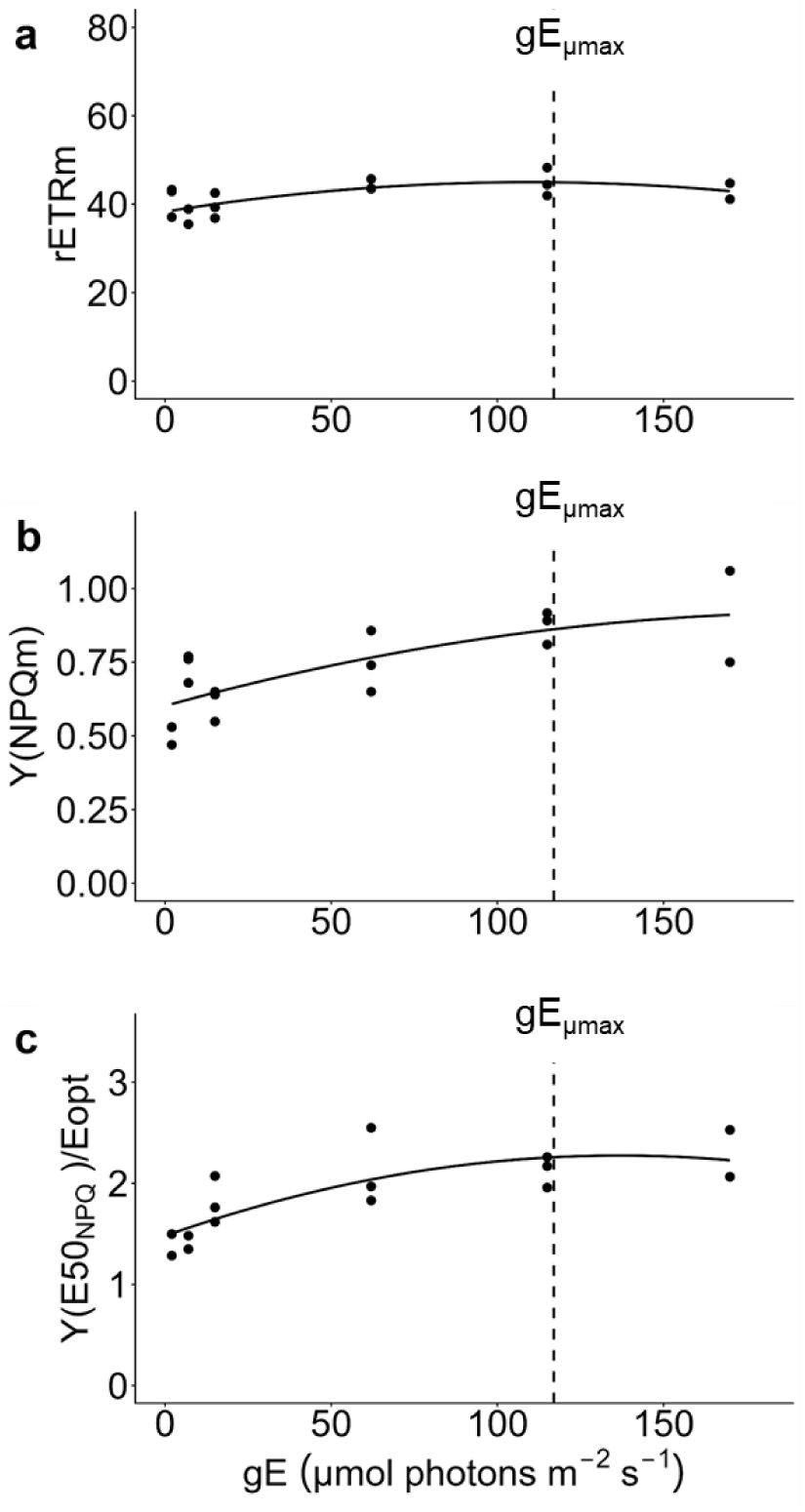
Photophysiological responses of *P. strigosum* to green-blue growth light intensity (gE). (**a**) rETRm, the maximum relative photosynthetic electron transport rate *versus* gE; (**b**) Y(NPQm), the quantum yield of maximal regulated non-photochemical quenching *versus* gE; (**c**) Y(E50_NPQ_) /Eopt, the ratio of the light intensity for which Y(NPQ) reaches half-maximal saturation to the optimal light intensity for photosynthesis *versus* gE. Dots are raw data, curves indicate regression models (second-order polynomial for (**a**) (R² = 0.47), (**c**) (R² = 0.57) and (**e**) (R² = 0.63) fitted to raw data. Dashed lines indicate gE = K_E_ (47 µmol photons m^-2^ s^-1^, the light saturation intensity for growth) and gE = gE_µmax_ (117 µmol photons m^-2^ s^-1^, the light at which the growth rate is maximal).

To better account for photoacclimation relative to the growth light regime, the ratios Ek/gE, Eopt/gE and Y(E50_NPQ_)/gE were examined as a function of gE (**Figure 6**) and of gE_µmax_ (**Figure S9**). Similar as for MPB (**Figure 6**), all three followed a logarithmic regression with values declining sharply towards 1:1 with increasing gE. Interestingly, while the intensity for reaching the 1:1 ratio was shifted by 50-60 µmol photons m^-2^ s^-1^ for Ek/gE and Eopt/gE in comparison to MPB values (**Figure 6a, b**), *i.e*., 74 vs 26 µmol photons m^-2^ s^-1^ and 155 vs 96 µmol photons m^-2^ s^-1^, the pattern for Y(E50)_NPQ_/gE was very close to the one of MPB (**Figure 6c**). When normalized to gE_µmax_ (**Figure S9**), Ek/gE and Eopt/gE converged to the 1:1 ratio in a perfect mirror image on both of gE_µmax_, *i.e.,* 0.63 for Ek/gE and 1.38 for Eopt/gE (**Figure S9a, b**).

## 4 Discussion

Underwater light availability varies in space and time, both in intensity and spectral composition, directly influencing photosynthetic life. This study investigated the photophysiological strategies of the subtidal microphytobenthos (MPB) *in situ* and of a representative of this microalgal community, the large-sized benthic diatom *Pleurosigma strigosum*, in response to ecologically relevant variations of underwater light climate. By exposing *P. strigosum* monospecific cultures to a gradient of green-blue growth light intensities representative of subtidal light environment at the surface of sediment, we documented a high photophysiological plasticity, providing, for the first time, insight into how a benthic diatom, and to larger extent MPB, cope with the subtidal light climate. Altogether, our results highlight the nature of subtidal photoadaptation: a very low green-blue light sensitive response, which likely underpin the ecological success of MPB in colonizing coastal subtidal sediments.

### 4.1 Photophysiology of the subtidal MPB community *in situ*

The subtidal MPB biomass maintains all year-around, especially during winter period (November-February) when the irradiance at the surface of sediment is the lowest (below 10 µmol photons m^-2^ s^-1^ and as low as 0.19 µmol photons m^-2^ s^-1^, **Figure 3**) and temperature is about 8-9°C (Chatterjee et al., 2013; Leynaert et al., 2018). MPB biomass first peaks in early spring (end of March-beginning of April), and shows secondary lower maxima later in Spring, Summer and early Autumn. While the first biomass maximum is strongly linked to increasing light availability and temperature (Chatterjee et al., 2013), secondary maxima are also influenced by nutrients availability (Laruelle et al., 2009; Chatterjee et al., 2013). In our study, we focused on deciphering the role of light only.

The photophysiological parameters rETRm, Ek, and Eopt followed a seasonal unimodal pattern closely tracking PAR at the surface of sediment and explaining half of their variability (48 to 55%, **Figure 5a, b**; **Figure S5b, d**). Such pattern mirrors previous reports on both subtidal (Light & Beardall, 2001) and intertidal MPB (Pniewski et al., 2015) and illustrates the tight coupling between ambient light, photoacclimation state (Ek, Eopt) and photosynthetic productivity (rETRm). The photoacclimative plasticity and robustness of subtidal MPB was further supported by a steady (max. 20% variation) Eopt/Ek ratio of 2.5 throughout the year, signifying permanently optimized light harvesting and light use efficiency relative to the prevailing *in situ* light climate (Lacour et al., 2017). The next striking feature of subtidal MPB photophysiology is its ability to maintain rETRm photosynthetic productivity (albeit at a level 15 times lower than its maximum) at irradiances close to 0 µmol photons m^-2^ s^-1^, and with Ek and Eopt as low as 4.5 and 9 µmol photons m^-2^ s^-1^, respectively (**Figure S5**). The very low light adaptation of subtidal MPB was further supported by the perfect match (ratio 1:1) of Ek/PAR versus *in situ* irradiance for a low value of 26 µmol photons m^-2^ s^-1^ (**Figure 6a**), indicating that the light saturation of MPB photochemistry was reached well before the maximum recorded PAR (6.5x higher). Similar very low light adaptation was previously reported in polar ecosystems where sea-ice sympagic and sub-ice planktonic diatom communities thrive under extremely low light (Hancke et al., 2018; Randelhoff et al., 2020; Hoppe et al., 2024) (see Section 4.4).

Oppositely, rETRm, Ek and Eopt reached their highest values from May to September (with a concomitant maximum by the end of June), likely illustrating a positive role of higher temperatures complementary to higher light availability (Chatterjee et al., 2013). All three parameters showed a maximum for a 100-120 µmol photons m^-2^ s^-1^ *in situ* PAR (**Table S2**) which strikingly coincides with the early Spring exponential increase in MPB biomass, 10-15 days before it reaches its maximum (Chatterjee et al., 2013). The reaching of PAR values ≥ 100 µmol photons m^-2^ s^-1^ was observed between mid-march to mid-september (**Figure 3**), covering the highest MPB biomass period. This PAR value thus appears to be a threshold for both subtidal MPB early Spring blooming (see also Chatterjee et al., (2013)) and photosynthetic productivity maintenance throughout the productive season. This threshold was further supported by two observations: (i) the light intensity for reaching rETRm perfectly matches the *in situ* irradiance close to 100 µmol photons m^-2^ s^-1^ (*i.e.,* Eopt/PAR *vs.* PAR = 1:1 for a PAR of 96 µmol photons m^-2^ s^-1^, **Figure 6b**); (ii) the ratio of intensities for the induction of photoprotection (Y(E50_NPQ_)) and optimal photochemistry (Eopt) reaches its lowest value (closest to 1) for a PAR of 100 µmol photons m^-2^ s^-1^ (**Figure 5f**).

Contrastingly to photochemistry, the photoprotection ability of subtidal MPB (Y(NPQm)) and its irradiance-induction (Y(E50_NPQ_)) were mostly stable across seasons and *in situ* PAR range. Mean Y(E50_NPQ_) was strikingly well above the maximum *in situ* PAR recorded (2.5x higher), which was also well illustrated by the consistently above 1 relationship of Y(E50_NPQ_)/PAR *vs.* PAR (**Figure 6e**). The fact that the light intensity needed to induce half NPQm always exceeds ambient PAR, highlights that (i) MPB photosynthetic productivity operates largely below its photoprotective potential under subtidal light conditions, (ii) even if unnecessary most of the year, NPQ potential is kept high, especially maintaining half of its maximum in winter, and can reach close to maximum (0.92 over 1 in April during first blooming, **Figure 5c**) values when punctually needed. The contrasting inverse seasonal tuning of Y(E50_NPQ_)/Eopt further shows that, during the cold (8-9°C, Leynaert et al., (2018)) dim winter months, subtidal MPB maintains a wide photoprotective margin induction *versus* photochemistry, which narrows during brighter spring and summer periods, reflecting a finely tuned balance between light harvesting and utilization, and limitation and prevention of photodamage. This ability was further supported by the sharp logarithmic decrease/increase during winter/spring and autumn/winter transitions of Y(E50_NPQ_)/Eopt over a very narrow range of very low *in situ* irradiances (0-10 µmol photons m^-2^ s^-1^). This behaviour matches the one depicted by Lacour et al., (2020) whereby, in contrast to other microalgal groups, diatoms are able to ensure efficient chloroplast bioenergetics thanks to maintaining a large photoprotective ‘buffer’ across a broad range of irradiances, temperatures and nutrient availability. This capacity especially allows very low-light adapted diatom species and communities, such as subtidal MPB and sea-ice sympagic diatoms, to (i) maintain effective photosynthetic machinery under low temperatures and/or irradiances close to 0 µmol photons m^-2^ s^-1^, and even prolonged darkness (Sundbäck & Jönsson, 1988; Randelhoff et al., 2020; Joli et al., 2024), (ii) rapidly enhance photochemical efficiency to support cell growth upon exponential irradiance increase by the end of winter/beginning of spring (Wulff et al., 2008; Veuger & Van Oevelen, 2011; Chatterjee et al., 2013; Kvernvik et al., 2018; Hoppe, 2022). This is the pattern we observe and which supports the blooming of subtidal MPB earlier in the season (Chatterjee et al., 2013), a timing of crucial importance for the bay of Brest food web (Longphuirt et al., 2007; Chatterjee et al., 2013).

### 4.2 Strong similarity between the subtidal benthic diatom *Pleurosigma strigosum* and the *in situ* subtidal MPB photophysiologies

When we grew subtidal MPB diatom *P. strigosum* under a light climate as close as possible to the *in situ* one, including irradiance range and light spectrum, we observed strong similarities between monospecific cultures and MPB community light-responses (Section 4.1). Especially, paramount rETRm and Y(E50_NPQ_)/Eopt parameters reached their maxima at similar growth intensities (gE) than *in situ* irradiance for MPB (100 and 137 µmol photons m^-2^ s^-1^, respectively) (**Table S2**). Working on an isolated representative of subtidal MPB allowed us to characterise the light-dependence of additional crucial photophysiological parameters which maxima were strikingly close to the *in situ* 100-120 µmol photons m^-2^ s^-1^ threshold (Section 4.1) in that order: pigments (82 ± 3 µmol photons m^-2^ s^-1^) < C:N (97 µmol photons m^-2^ s^-1^) < daily C production (98 µmol photons m^-2^ s^-1^) < Chl *a*:C (110 µmol photons m^-2^ s^-1^) < µmax (117 µmol photons m^-2^ s^-1^) (**Table S2**), furthermore confirming the crucial significance of this irradiance level in subtidal MPB diatom photobiology. This is consistent with the ≥ 100-250 µmol photons m^-2^ s^-1^ photobiology threshold often reported for higher light-adapted intertidal MPB and epileptic diatoms depending on seasons and species (see Barnett et al., (2020) and references therein).

Interestingly, while rETRm *P. strigosum* followed a similar irradiance-dependent pattern as the one of MPB (**Figures 9a, 5b**), it was not the case of Ek and Eopt that were stable across the gE gradient and in the upper range of MPB values. It was the opposite for Y(E50_NPQ_): stable in MPB, increasing with gE in *P. strigosum*, and reaching its maximum value (similar to Y(E50_NPQ_) MPB) for the highest gE. Also, the ratios of light optima over gE (Ek/gE, Eopt/gE, Y(E50_NPQ_)/gE) *vs.* gE followed the same trend as MPB ratios over *in situ* PAR (**Figure 6**) albeit a shift towards higher intensities (+50 µmol photons m^-2^ s^-1^ for Ek and Eopt). These discrepancies are probably due to laboratory optimal temperature (*i.e.,* 15-16°C summer-like *in situ* subtidal level, Leynaert et al., (2018)) and nutrient replete growth conditions, comparatively to *in situ*, which leverage part of the excitation pressure on the photosynthetic apparatus in *P. strigosum*. This is furthermore illustrated by its higher rETRm compared to MPB, and the high (*i.e*., in the upper range of MPB values) and stable Fv/Fm across gEs (**Figure 7a**), supporting no apparent and permanent photodamage in *P. strigosum* (Frankenbach et al., 2018). Nevertheless, and as for MPB, the Eopt/Ek ratio was stable across gEs, and most importantly similar to MPB mean value (2.25 *vs*. 2.53), and Y(E50_NPQ_)/gE *vs.* gE consistently remained higher than gE (**Figure 6e**). Altogether, these patterns indicate that, both in *P. strigosum* and *in situ* MPB, photochemical light intensity optima (Ek and Eopt) concomitantly converged towards the prevailing irradiance over a reduced range of intensities representative of *in situ* subtidal light climate (*i.e.,* up to 170 µmol photons m^-2^ s^-1^), while Y(E50_NPQ_) always remained well above gE and PAR, respectively.

Altogether, these common and differential features also point out the main difference between our experimental light climate and the *in situ* one: in our laboratory conditions *P. strigosum* acclimated to prolonged (several weeks) continuous light levels, when *in situ* MPB is exposed to regular changes in incident irradiance due to the two-week tidal cycle. While problematic in appearance, these differences are very much informative. When the tidal coefficient is strong, weather is sunny and tide is the lowest, the irradiance at the surface of sediment surely exceeds the maximum measured here and elsewhere (*ca.* 200 µmol photons m^-2^ s^-1^, (Chatterjee et al., 2013)). The adaptive strategy of MPB is a seasonal dynamic concomitant (stable Ek/Eopt) modulation of light optima Ek and Eopt strictly connected to PAR (*i.e.,* Ek/PAR and Eopt/PAR *vs.* PAR patterns, **Figure 6**) and with a strong modulation of the relationship between photochemistry and photoprotection (Y(E50_NPQ_)/Eopt). This strategy is obviously central as also observed in *P. strigosum* (similar Ek/Eopt, Ek/gE and Eopt/gE *vs.* gE patterns, **Figure 6**). In parallel, MPB maintains Y(E50_NPQ_) and Y(NPQm) high and stable, while keeping its potential for enhancing Y(NPQm) to a maximum (*i.e.,* close to 1). This is further strengthened by the ability of *P. strigosum* to increase Y(NPQm) and Y(E50_NPQ_) up to the maximal values recorded in MPB. This strategy allows to maintain Fv/Fm fairly high and stable and to increase rETRm with seasonal irradiance. The high Y(E50_NPQ_) and Y(E50_NPQ_)/PAR *vs.* PAR pattern indeed allow MPB not to induce NPQ for too low irradiances and thus keeps the potential for photochemistry optimal for the dominant irradiance range at the surface of sediment (*i.e.*, 0 to 200 µmol photons m^-2^ s^-1^), similar as reported before in planktonic diatoms (Lavaud et al., 2007). High and stable Y(E50_NPQ_) makes full sense during the strongest irradiance period (April-September, **Figure 5e, f**) when Eopt moves to higher irradiances, to fit the seasonal light demand, getting closer to Y(E50_NPQ_) (*i.e.,* Y(E50_NPQ_)/Eopt approaching 1, **Figure 5e**) without reaching it. This situation is further strengthened by the similar Y(E50_NPQ_)/Eopt (*ca.* 2, **Figure 9e**) at the highest gE in *P. strigosum*. Therefore, Y(E50_NPQ_) seems to work as an essential ‘safety margin’ for modulating the photochemical potential over a broad range of seasonal and experimental prolonged light exposures and more punctual tidal-driven higher light exposures. All and all, this is a remarkable illustration of a high photophysiological plasticity modelled together on the long-term seasonal (days and weeks) irradiance changes (for both photochemistry and photoprotection capacities) and on a shorter time-range (hours) tidal cycle-driven light fluctuation (for photoprotection capacity).

Contrary to *in situ*, in our experimental set-up, the photoperiod was unchanged (16 h L:8 h D) and the subtidal diel photoperiodic vertical migration (Longphuirt et al., 2006) of *P. strigosum* was prevented, so that the regulation of photosynthetic productivity only relied on photoacclimation to increasing light doses (*i.e*., 0.12 mol photons m^-2^ d^-1^ to 9.8 mol photons m^-2^ d^-1^). The similarity between *P. strigosum* and MPB thus strongly suggests that the photophysiological response alone can sustain photosynthetic efficiency under the subtidal light climate, highlighting the adaptive value of intrinsic photophysiological flexibility and robustness in benthic diatoms, even more in non-motile epipsammic forms (Jesus et al., 2009; Cartaxana et al., 2011; Barnett et al., 2015; Blommaert et al., 2017). It also confirms that vertical migration is most probably mainly used by subtidal MPB diatoms to access light at the surface of sediment (Longphuirt et al., 2006) instead of being used as a photoprotective behaviour (*i.e*., negative phototaxis). This is in marked contrast to intertidal MPB where epipelic motile diatoms also use vertical migration for positioning themselves along the sharp sediment vertical light gradient and for escaping from low tide excess light exposure at the surface of sediment (Consalvey et al., 2004; Jesus et al., 2023; Serôdio et al., 2023).

Conclusively, and although *P. strigosum* is not a dominant component of subtidal MPB in the bay of Brest, the general coherence between its light-response and the one of *in situ* MPB under a similar light climate offers a unique model species and growth form to further decipher the specific light response of subtidal MPB under precisely controlled light conditions.

### 4.3 Photophysiological plasticity of subtidal *Pleurosigma strigosum*

In the bay of Brest, at *ca.* 10 m depth, subtidal MPB diatoms are exposed year-round to relatively weak irradiances, rarely approaching/exceeding 200 µmol photons m⁻² s⁻¹ at the sediment surface at high tide, and with a spectral composition strongly shifted toward green-blue wavelengths (**Figures 3, S3**; (Chatterjee et al., 2013; Leynaert et al., 2018)). In order to better characterise the photophysiology of *P. strigosum*, we grew it under a range of light intensities (gE) in accordance with seasonal *in situ* irradiances, *i.e.,* from 2 to 170 µmol photons m^-2^ s^-1^.

All elementary composition parameters, pigment contents and photophysiological traits exhibited a strict light-dependent hyperbolic relationship with gE with two crucial thresholds: K_E_ (47 µmol photons m^-2^ s^-1^) and gE_µmax_ (117 µmol photons m^-2^ s^-1^) (see **Figures 7, 8, 9**). Such pattern reflects the progressive transition from light-limited to light-saturated photosynthetic regimes, whereby cells modulate their pigment content, photosynthetic capacity, and photoprotective energy dissipation pathways to maintain photochemical and metabolic balance under varying irradiance levels. While this is generally consistent with classical models of photoacclimation and photosynthesis *vs.* irradiance (MacIntyre et al., 2002), and with the unique and central role of NPQ regulation in diatom photophysiology and growth strategy (Lacour et al., 2020), there were some unusual features pointing out to a unique adaptability to (very) low green-blue irradiances while retaining a high ability for photoprotection, allowing for a precise tuning of light harvesting and utilisation to reach an optimal match between prevailing light conditions, C fixation and growth (see **Figures S6b, S10**).

While the *P. strigosum* photochemical and NPQ parameters were generally consistent with the ones of *in situ* MPB (see Section 4.2), a further examination brought additional evidence on the tight link between photochemistry and growth, and on the central role of NPQ. rETRm increased and saturated with gE, when NPQ rose steadily, reflecting the progressive engagement of photoprotective pathways (**Figures 9a, b**). While up to half gE_µmax_, NPQ increase can be attributed to the nearly doubling of the DD+DT cell content (**Figure 6b**), the further increase under higher intensities was likely supported by the synthesis of LHCx proteins (Buck et al., 2019; Croteau et al., 2025). When the light optima for photochemistry Ek and Eopt remained stable, and high, across the light gradient, the irradiance threshold for NPQ induction (Y(E50_NPQ_)) increased linearly with gE (**Figure S8**), indicating a strong adjustment of photoprotection dynamics to incident irradiance. Interestingly, the mean Eopt/Ek ratio was very close to the one of gE_µmax_/K_E_ (2.25 *vs.* 2.49) indicating a strong link between photochemistry and growth light-limited and growth light-saturated thresholds. When normalized to gE_µmax_, the convergence of light intensity optima at the equidistance of gE_µmax_ (*i.e.,* 0.63 for Ek/gE and 1.38 for Eopt/gE) (**Figure S9**) underscores a strong adaptive tuning between photochemistry and growth rate as a function of prevailing irradiance. This is furthermore illustrated by the powerful relationship between daily C production, C cell content, Chl *a*:C and C:N *versus* growth rate relationships (**Figure S10**). Conversely, the consistent maintenance of Y(E50_NPQ_)/gE well above gE_µmax_ (**Figures 6, S9**) further supports the ‘safety margin’ role proposed above (see Section 4.2).

The most remarkable feature of *P. strigosum* light-response was the light-dependent hyperbolic pattern of C-related parameters with increasing gE and over a relatively restricted gE range. While the cell C content showed an expected logarithmic decrease with increasing gE (**Figure S6a**) due to enhanced investment in growth (*i.e.,* opposite to C storage along with very low growth rate under very low intensities; (Lacour et al., 2018, 2022); see **Figure S6b**), C:N and Chl *a*:C showed completely different patterns as compared to the diatom textbook ones (MacIntyre et al., 2002; Lacour et al., 2017). Typically, Chl *a*:C shows logarithmic decrease with increasing irradiance due to the concomitant reduction of the cell investment in Chl *a* synthesis and light-harvesting proteins beyond the light-saturating threshold, and the opposite under light-limiting conditions. According to MacIntyre et al., (2002), it reflects a trade-off between C allocation to the photosynthetic *versus* biosynthetic machinery, maintaining photochemistry near the point of light saturation. This trade-off is further illustrated by the stable C:N ratio across irradiance supporting fine-tuned metabolism for a stable nutrient allocation (MacIntyre et al., 2002; Lacour et al., 2017, 2018). In *P. strigosum*, both C:N and Chl *a*:C showed a second-order polynomial relationship with gE, increasing up to K_E_, reaching a maximum up to gE_µmax_ and decreasing above gE_µmax_ (**Figures 7b, c**). The plateauing of Chl *a*:C and further decrease from K_E_ on is expected, *i.e.,* the growth rate and daily C production become light-saturated and the cells synthesise less Chl *a* and other light-harvesting pigments (especially Fx and Chl *c*, **Figure 8a**) to reduce the light energy absorption (see **Figure S10**). The Chl *a*:C increase up to K_E_ is totally unusual. To our best knowledge, such pattern was reported only in (very) low light- and cold-adapted arctic diatoms, and in the planktonic *Skeletonema costatum* (Falkowski & Owens, 1980), although for a twice lower range of intensities (*i.e.,* maximal Chl *a*:C for 50 µmol photons m^-2^ s^-1^) (see Lacour et al., (2018); Croteau et al., (2022)). We hypothesise that such a pattern is based on the biofilm-living form of benthic diatoms at the surface of sediment (and which we also observe in our *P. strigosum* cultures), a collective structuring which decreases even more the light accessibility through self-shading. The increase in incident irradiance being much attenuated by the cell stacking, they linearly accumulate Chl *a* and other pigments to enhance their light-harvesting capacity up to a certain threshold (*i.e.,* K_E_), and thereafter, from K_E_ to before gE_µmax_, cells stop accumulating pigments. As a consequence of enhancing light-harvesting capacity, the daily C production increases, supporting higher growth rate and concomitant expected decrease in the C cell content, and explaining the unusual increase and plateauing of Chl *a*:C and C:N ratios. Another process that exists only in large-sized diatoms, such as *P. strigosum*, and which can add a layer of fine-tuning at a spatial scale intermediate between the biofilm and the light-harvesting antenna, is the karyostrophy, *i.e.,* the light intensity-driven plastid contraction (Bastos et al., 2025). By contracting the plastids around the central nodule, karyostrophy generates light attenuation in the proximal region of *P. strigosum* cells and could very well control the light accessibility within the biofilm. Strikingly, it was shown to be significantly induced from 60 µmol photons m^-2^ s^-1^ on (Bastos et al., 2025), just above K_E_, thus possibly being involved in framing the paramount K_E_-gE_µmax_ light range.

### 4.4 Subtidal *Pleurosigma strigosum* shows photophysiological features close to the very low-light adapted polar diatoms

Surprisingly, and even if the growth temperature was similar to *in situ* subtidal summer-like level (*i.e.,* 15-16°C, (Leynaert et al., 2018)), *P. strigosum* retains obvious features of (very) low light- and cold-adapted diatoms, positioning its photophysiology closer to the polar diatoms than their temperate counterparts (Lacour et al., 2017).

In Sections 4.1 and 4.3, we discussed how the maintenance of a high NPQ potential allows subtidal MPB and polar diatoms to preserve an effective photosynthetic machinery under low temperatures and/or irradiances close to 0 µmol photons m^-2^ s^-1^, and even prolonged darkness, a crucial ecophysiological feature in diatom growth strategy that supports their success in harsh habitats characterised by extreme light climates (Lacour et al., 2020). The predicted NPQm of 0.60 for a gE of 0 µmol photons m^-2^ s^-1^ (**Table S5**), which fits well with the lowest winter MPB NPQm values (**Figure 5c**), further confirms this assumption. Along with the conserved high NPQ potential at 0 µmol photons m^-2^ s^-1^, *P. strigosum* exhibited a high light-harvesting capacity (*i.e.*, high Chl *a*, Fx and Chl *c* cell contents) and a high potential for photochemistry (rETRm) (**Table S5**). Furthermore, Eopt/Ek was 2 and Y(E50_NPQ_)/Eopt was close to 1 (**Table S5**), confirming the ability of *P. strigosum* to maintain, together with an effective photosynthetic machinery, a nearly optimal relationship between photochemistry thresholds and photoprotection induction, a feature essential for the immediate reset of photosynthesis upon light re-exposure after prolonged darkness (see Section 4.1). The advantage of keeping NPQ high under limiting gE rather than degrading Chl *a*, other pigments, and part of the photosynthetic machinery (Morin et al., 2020; Joli et al., 2024) could be to maximise light harvesting at very low intensities during the winter dim period, and with the additional extreme attenuation within the cell biofilm structure (see Section 4.3). Such a regulation would explain the unusually very low Chl *a*:C values (9 to 16 mg g^-1^) at gE 2-7 µmol photons m^-2^ s^-1^, even compared to the lowest value of 22 mg g^-1^ for similar lowest gE in Arctic diatoms (see Croteau et al., (2022)). Furthermore, because it takes several weeks to reach a PAR close to gE_µmax_, and MPB Eopt, from the end of winter to the beginning of spring (**Figure 3**), it is necessary for *P. strigosum*, one of the first bloomers (Leynaert et al., 2018), to exploit at best daily periods of low light (Lacour et al., 2020). This behaviour very much resembles the one of sea-ice sympagic diatoms when getting out of the polar night (Croteau et al., 2022; Guérin et al., 2024).

Another paramount parameter to consider is the C:N ratio, which is a key ecophysiological indicator of nutrient allocation and metabolic status (Lacour et al., 2017). In general, it was low (4.8 ± 1.4) and predicted 2.45 at 0 µmol photons m^-2^ s^-1^ (**Table S5**). Such elementary composition is further consistent with low-light and/or cold-adapted diatoms, where enhanced N investment in photosynthetic and biosynthetic machinery supports efficient resource use under limiting light energy supply (Lacour et al., 2017, 2022). In polar diatoms, low C:N (typically below the 6.6 Redfield ratio) is explained by a high Rubisco content that alleviates the slow enzyme rate due to low temperatures (Young et al., 2015; Young & Schmidt, 2020; Lacour et al., 2022). It is possible that *P. strigosum*, due to its large biovolume (*ca.* 30 000 µm^3^), retains a large quantity of Rubisco (*i.e.,* larger than small-sized temperate benthic diatoms), that would explain the high C cell content at (very) low intensities, even compared to Arctic diatoms (0.50 ± 0.05 pg C µm^-3^ *vs.* 0.27 ± 0.05 pg C µm^-3^, **Figure S6a** and Lacour et al., (2022)), and further explains the maintenance of a 0 µmol photons m^-2^ s^-1^ predicted Chl *a*:C (17.5 mg g^-1^, **Table S5**) close to the one of Arctic diatoms (20-25 mg g^-1^; Croteau et al., (2022)). Possibly, this hypothetical higher Rubisco content would help *P. strigosum* to support daily C production during the dim and cold winter period.

The further examination of growth (µmax = 0.37 d^-1^), elementary composition (C:N = 4.8 ± 1.4), and photoadaptive (K_E_ = 47 µmol photons m^-2^ s^-1^, α_µ_ = 0.91 d^-1^ m^-2^ mol photons) features showed an alignment with the mid/upper range of polar diatoms values and the lower/lowest range of temperate diatoms (MacIntyre et al., 2002; Sarthou et al., 2005; Lacour et al., 2017). More specifically, the general growth- and light-response of *P. strigosum* fairly matches the one of the typical Arctic sea-ice edge species *Thalassiosira gravida* (Lacour et al., 2018; Croteau et al., 2022) with close µmax (0.37 *vs.* 0.32 d^-1^), gE_µmax_ (117 *vs.* 88 µmol photons m^-2^ s^-1^), with a light efficiency use α_µ_ of 0.91 *vs*. up to 0.88 d^-1^ m^-2^ mol photons in *T. gravida*, and with similar ranges of Chl *a*:C values (*ca.* 10 to 50 *vs.* 10 to 65) and max C:N ratio (6.3 *vs.* 6.5). This is noticeable as *T. gravida* was categorised as using a ‘layer former’ strategy (oppositely to a ‘mixer’ strategy), illustrated by an adaptation to (very) low and relatively stable (*i.e.,* reduced and low frequency light fluctuations) irradiances (see Lacour et al., (2018) and references therein). Noteworthy, the *in situ* subtidal light climate of *P. strigosum* is very similar to the one of Arctic diatoms during the productive end of winter-to-beginning of summer period with a light dose increasing from nearly 0 to *ca.* 10 mol photons m^-2^ d^-1^ and a light spectrum strongly shifted toward green-blue wavelengths (Leynaert et al., 2018; Guérin et al., 2024).

## 5 Conclusion

Understanding how microphytobenthos (MPB) diatoms respond to their light environment is key to assessing their productivity in sediments and their ecological role in coastal estuarine ecosystems. This field of research has become even more crucial with the sea-level rise and the progressive submersion of intertidal sediments (Murray et al., 2019) which generates a profound change in the light climate with the conversion to a subtidal mode, with dramatic consequences on MPB productivity (Mangan et al., 2020; Flowers et al., 2023). Therefore, subtidal MPB is an open window to the future of present intertidal MPB. This study investigated the photoadaptive strategy of subtidal (*ca.* 10 m depth) MPB in the semi-enclosed bay of Brest, and of one of its representatives: the large-sized motile diatom *Pleurosigma strigosum*. Our results indicate a unique coordinated photoadaptive strategy between photochemistry, photoprotection and resource allocation, consistent with a subtidal lifestyle, where non-photochemical quenching-NPQ plays a central role in fine-tuning the cells bioenergetics, their productivity and ultimately their capacity (i) to thrive under (very) low-light subtidal conditions, (ii) to bloom early in the season as soon as the subtidal irradiance at the surface of sediment reaches *ca*. 100 µmol photons m^-2^ s^-1^, *i.e.*, 20x lower than full sunlight. The general coherence between subtidal *P. strigosum* and MPB light-responses offers a unique model species and growth form to further decipher the specific light-driven metabolism of subtidal MPB under precisely controlled environmental conditions. In the present study, we focused on the response of *P. strigosum* to a range of low green-blue light intensities reproducing the subtidal light climate at high tide. It would be of interest to further decipher two paramount photoadaptive features of subtidal *P. strigosum*: (i) its ability to respond to higher punctual light exposures representative of low tide periods to verify if it retained photoprotective and recovery abilities closer to very low-light adapted growth forms (Croteau et al., 2021, 2026) or to higher light adapted intertidal benthic diatoms (Barnett et al., 2015; Blommaert et al., 2017); (ii) its wavelength-specific response based on the 10 m light spectrum that is strongly shifted towards green-blue and nearly deprived in red radiations.

## Acknowledgements

The authors gratefully acknowledge the technical support of the analytical platform PACHIDERM for BSi and POC/PON analyses, Beatriz Beker for her help with identifying the species *Pleurosigma strigosum*, Ingrid Kernalléguen for her help with the MEB pictures, Thomas Lacour for his help in interpreting Carbon data, and Julien Thébault for kindly providing the map of the bay of Brest.

## Data availability statement

data can be found on SEANOE (DOI: 10.17882/111840).

## Acknowledgements, including funding statement and permission to reproduce material from other sources

The author(s) declare that financial support was received for the research, authorship, and/or publication of this article. This work was supported by the Région Bretagne and the Doctoral School ‘Sciences de la Mer et du Littoral’ (PhD fellowship to Anna Isaia), the French CNRS EC2CO Microbiome program (project LUMIÈRE), the EUR ISblue-Interdisciplinary graduate school for the blue planet (ANR-17-EURE-0015) co-funded by the French government under the France 2030 program ‘Investissements d’Avenir’ (projects DIATOP and RAMI), the UBO/SEA-EU research program co-funded by the European Union and France 2030 (project RAMI), the French Ministry for Europe and Foreign Affairs, and the French Ministry for Higher Education and Research, the EU Horizon Research and Innovation Actions (project REWRITE, Grant Agreement 101081357).

The authors gratefully acknowledge the technical support of the analytical platform PACHIDERM for BSi and POC/PON analyses, Beatriz Beker for her help with identifying the species *Pleurosigma strigosum*, Ingrid Kernalléguen for her help with the MEB pictures, Thomas Lacour for his help in analysing Carbon data, and Julien Thébault for kindly providing the map of the bay of Brest.

## Ethics statement

## Author contributions

