## Supplemental Figures and Tables for "The peculiar photobiology of a giant motile diatom inhabiting the subtidal sediments of the bay of Brest"

**Material and methods**

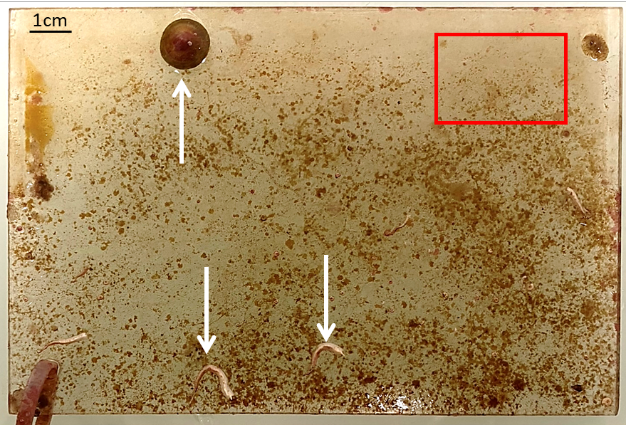

**Figure S1**: Typical example of a Plexiglas plate colonized by microphytobenthos and macrofauna at the Lanvéoc subtidal sediment (**Figure 1**). The red rectangle shows a dense biofilm zone, and white arrows point to benthic macrofauna.

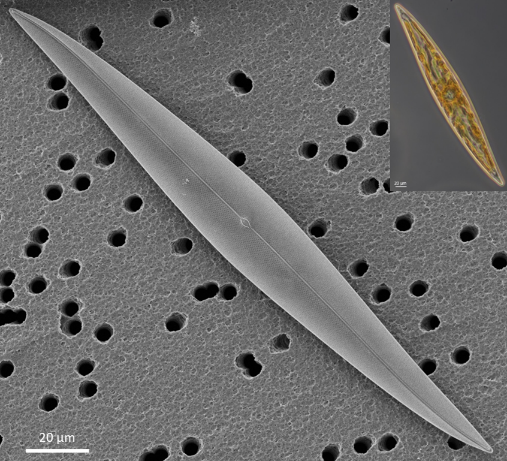

**Figure S2**: Electron and optical (top right) microscopic images of *Pleurosigma strigosum* isolated in the subtidal sediment of the Bay of Brest.

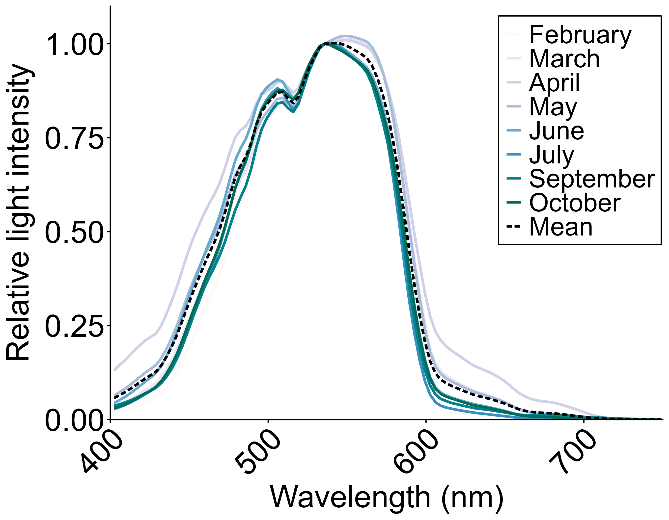

**Figure S3**: Seasonal variations in the emission spectrum measured at the sediment surface of the study site; the dashed black line represents the mean of all spectra used, as a model spectrum, for the green-blue culture conditions (see **Figure 2**).

**Table S1:** Definitions of parameters and units used in the study.

| Parameter | Definition | units |
| --- | --- | --- |
| *Growth performances, elemental composition* | |  |
| µ | Specific growth rate | day^-1^ |
| α_µ_ | growth rate | day^-1^ µmol photons m^-2^ s^-1^ |
| µmax | Maximal growth rate | day^-1^ |
| K_E_ | Light saturation intensity for growth | µmol photons m^-2^ s^-1^ |
| gE_µmax_ | Maximal growth light intensity | µmol photons m^-2^ s^-1^ |
| C | Carbon content | pg µm^-3^ |
| C : N | Carbon and Nitrogen content ratio | g g^-1^ |
| Chl *a* : C | Cellular chlorophyll *a* and Carbon ratio | mg g^-1^ |
| Daily C production | Daily Carbon production | ng cell^-1^day^-1^ |
| BSi | Biogenic Silica | nmol cell^-1^ and nmol m^-2^ |
| Chl *a* | Cellular chlorophyll *a* concentration | pg cell^-1^ and mg m^-2^ |
| *Pigment content* |  |  |
| Chl *c* | Chlorophyll *c* | pg cell^-1^ and mol 100 mol Chl *a*^-1^ |
| Fx | Fucoxanthin | pg cell^-1^ and mol 100 mol Chl *a*^-1^ |
| β-Car | β-Carotene | pg cell^-1^ and mol 100 mol Chl *a*^-1^ |
| DD + DT | Xantophyll cycle pigments: diadinoxanthin and diatoxanthin | pg cell^-1^ and mol 100 mol Chl *a*^-1^ |
| *Photosynthetic and photoprotection parameters* | |  |
| Fv | Variable fluorescence | rel. unit |
| Fm | Maximal fluorescence | rel. unit |
| Fv/Fm | Maximal quantum yield of photosytem II (PSII) photochemistry | rel. unit |
| rETRm | Maximal relative electron transport rate | rel. unit |
| Ek | Light saturation coefficient | µmol photons m^-2^ s^-1^ |
| Eopt | Light intensity for reaching rETRm | µmol photons m^-2^ s^-1^ |
| Y(NPQm) | Maximal regulated non-photochemical quenching yield | rel. unit |
| Y(E50_NPQ_) | Intensity for reaching 50% of Y(NPQm) | µmol photons m^-2^ s^-1^ |

**Results**

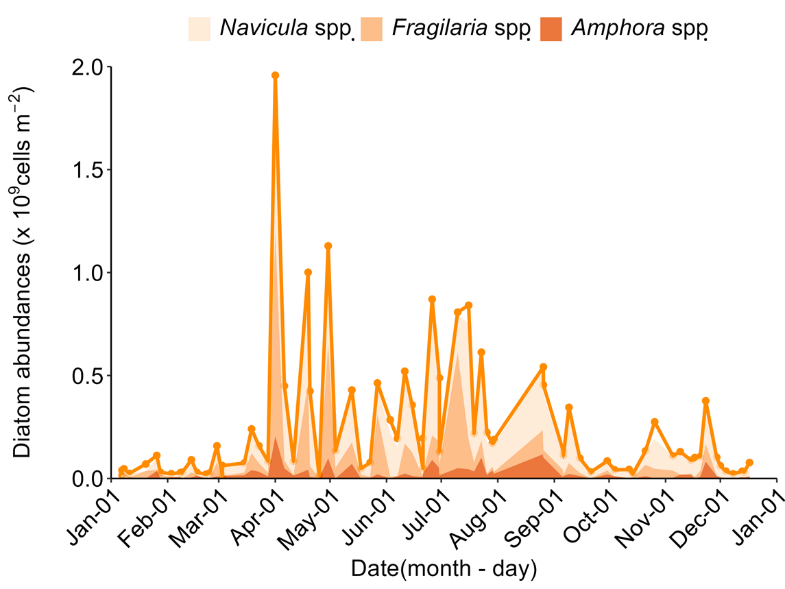

**Figure S4:** Annual profile of the total diatom abundance (orange dots and line) in the microphytobenthos at the Lanvéoc study site, as shown in **Figure 4**. The areas below the orange line show the detailed abundances for the three dominant diatom genera: *Navicula*, *Fragilaria* and *Amphora*.

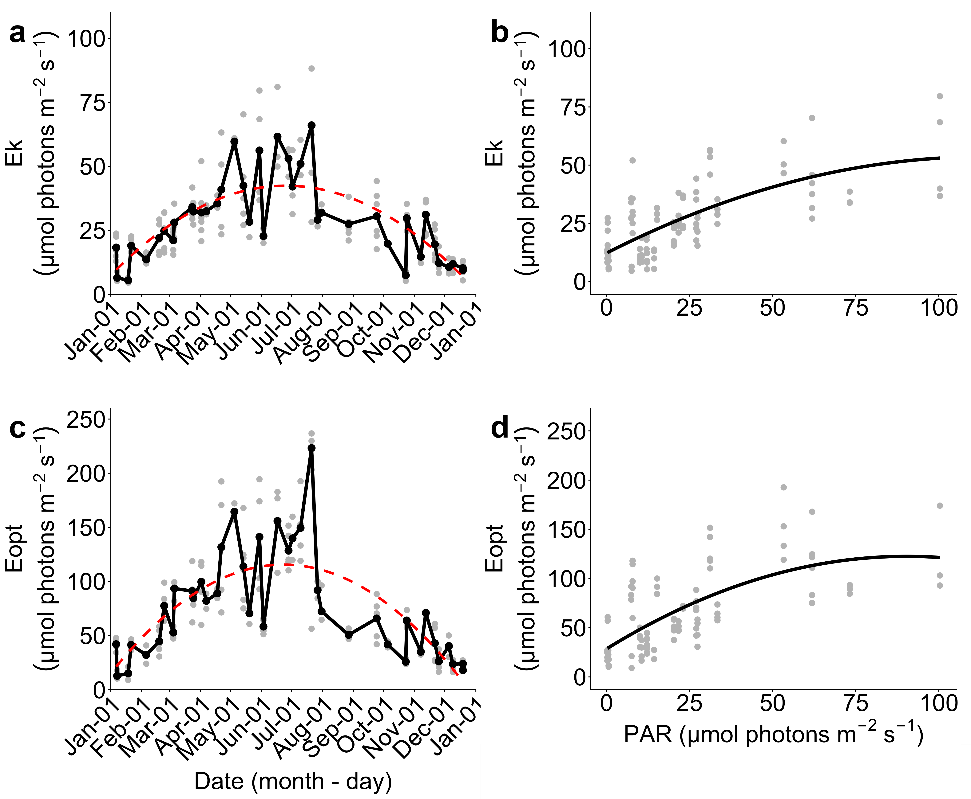

**Figure S5:** Annual dynamics of microphytobenthos photophysiological light use parameters (see **Table S1** for definitions) at the study site and their relationship with the corresponding photosynthetically active radiation (PAR) at the sediment surface. (**a**) Annual Ek; (**d**) Ek *versus* corresponding PAR (**c**) annual Eopt; and (**d**) Eopt *versus* corresponding PAR. Dots are raw data. Dashed red curves represent second-order polynomial regression fitted to raw data, R² are: (**a**) 0.51, (**b**) 0.52, (**c**) 0.55 and (**d**) 0.48.

**Table S2**: Intensity reached for the maximum or minimum (*) of each parameter from *in situ* microphytobenthos (MPB) (PARmax/min, photosynthetically active radiation for maximum or minimum) and *P. strigosum* grown under green-blue (GB) light (gEmax, growth light intensity for maximum) and the ratio of gEmax over gE_µmax_, the growth light intensity for maximal growth rate. Parameters were extracted from second-order polynomial and power regression from **Figures 5, 7, 8, 9, S5**, the definitions of the parameters are found in **Table S1**.

| *In situ* MPB | PARmax/min (µmol photons m^-2^ s^-1^) |  |
| --- | --- | --- |
| rETRm | 122 |  |
| Ek | 114 |  |
| Eopt | 113 |  |
| Y(E50_NPQ_)/Eopt* | 100 |  |
| GB *P. strigosum* cultures | gEmax (µmol photons m^-2^ s^-1^) | gEmax/gE_µmax_ |
| DD+DT (pg cell^-1^) | 79 | 0.67 |
| Chl *c* (pg cell^-1^) | 80 | 0.68 |
| Chl *a* (pg cell^-1^) | 82 | 0.70 |
| Fx (pg cell^-1^) | 82 | 0.70 |
| β-car (pg cell^-1^) | 85 | 0.73 |
| C : N | 97 | 0.83 |
| Daily C production (ng cell^-1^ day^-1^) | 98 | 0.84 |
| rETRm | 110 | 0.94 |
| Chl *a* : C | 110 | 0.94 |
| **µmax (day^-1^)** | **117** | **1.00** |
| Y(E50_NPQ_)/Eopt | 137 | 1.17 |
| Y(NPQm) | 204 | 1.74 |

**Table S3**: Results from *post hoc* (Wilcoxson rank sum test) tests of the maximum quantum efficiency of PSII (Fv/Fm, **Figure 5a**), and the frustule length and width (**Table S4**) of *P. strigosum* grown under increasing green-blue light intensities (gE). Differences were considered significant for *p-value* < 0.050 (bold values).

|  | gE | 2 | 7 | 15 | 62 | 115 |
| --- | --- | --- | --- | --- | --- | --- |
| Fv/Fm | 7 | 1.0000 |  |  |  |  |
|  | 15 | 1.0000 | 1.0000 |  |  |  |
|  | 62 | 1.0000 | 1.0000 | 1.0000 |  |  |
|  | 115 | 1.0000 | 0.3490 | 0.5120 | 0.0680 |  |
|  | 170 | 1.0000 | 1.0000 | 1.0000 | 1.0000 | 0.1660 |
|  |  | 2 | 7 | 15 | 62 | 115 |
| length | 7 | **1.50E-03** |  |  |  |  |
|  | 15 | **7.20E-03** | **3.90E-05** |  |  |  |
|  | 62 | **1.12E-02** | **3.90E-05** | 0.8198 |  |  |
|  | 115 | 0.8198 | **3.90E-05** | **1.90E-03** | **1.05E-02** |  |
|  | 170 | 0.1729 | **3.90E-05** | **1.10E-07** | **1.40E-07** | **1.80E-03** |
|  |  | 2 | 7 | 15 | 62 | 115 |
| width | 7 | 0.1471 |  |  |  |  |
|  | 15 | 1.0000 | 0.3237 |  |  |  |
|  | 62 | 0.1196 | **3.80E-03** | 0.3237 |  |  |
|  | 115 | 1.0000 | 1.0000 | 1.0000 | **3.00E-03** |  |
|  | 170 | 1.0000 | 0.2287 | 1.0000 | 1.0000 | 0.7237 |

**Table S4**: Frustule morphometrics, length and width of *P. strigosum* grown under increasing green-blue light intensities (gE). Data are expressed as mean ± SD. Statistical tests used were Kruskal-Wallis test and *post hoc* test (Wilcoxson rank sum test, see **Table S3**. Differences were considered significant for *p-value* < 0.050 (bold value), different letters indicate significant differences between light culture conditions.

| gE  (µmol photons m^-2^ s^-1^) | 2 | 7 | 15 | 62 | 115 | 170 | *p value* |
| --- | --- | --- | --- | --- | --- | --- | --- |
| **Frustule length (µm)** | 199.9 ± 5.9^ad^ | 178.8 ± 5.0^b^ | 205.3 ± 2.8^c^ | 205.0 ± 2.8^c^ | 201.8 ± 2.8^a^ | 198.2 ± 2.3^d^ | **8.93E-15** |
| **Frustule width (µm)** | 23.8 ± 0.5 | 22.7 ± 1.3^a^ | 24.0 ± 1.2 | 24.8 ± 1.0^b^ | 23.5 ± 0.8^a^ | 24.2 ± 1.2 | **6.99E-04** |

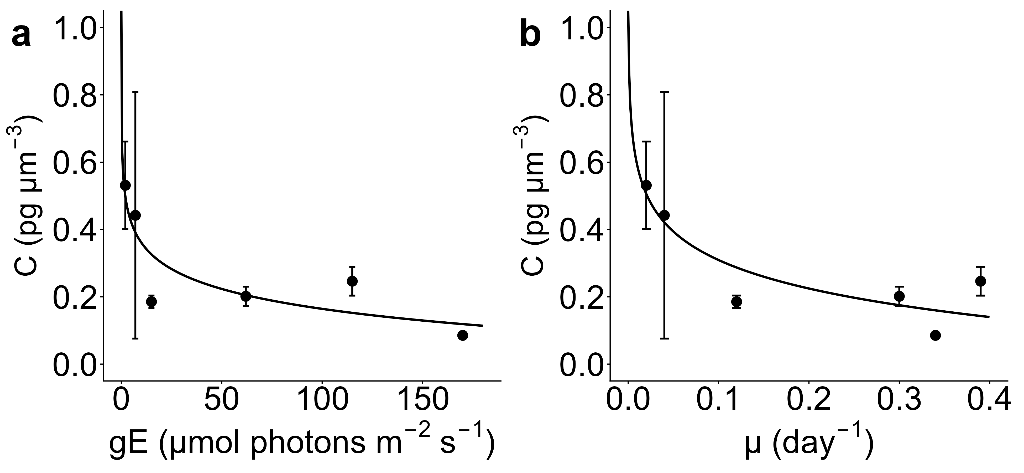

**Figure S6**: Cell C content of *P. strigosum* grown under increasing green-blue light intensities (gE): (**a**) as a function of gE, (**b**) as a function of growth rate (see **Figure 7a**). Grey dots are raw data; and black dots are mean. Solid lines are (**a**) logarithmic regressions with (**a**) R² = 0.77 and (**b**) R² = 0.80. The cell *P. strigosum* volume was estimated as 32330 µm^3^.

**Table S5**: *P. strigosum* parameter’s values predicted from fits (see **Figures 7, 8, 9, S6, S8**) for a green-blue growth light intensity of 0 µmol photons m^-2^ s^-1^. See **Table S1** for parameters’ definition.

|  | Parameter’s value |
| --- | --- |
| µmax (day^-1^) | 0 |
| Chl *a* (pg cell^-1^) | 126.6 |
| Fx (pg cell^-1^) | 78.3 |
| Chl *c* (pg cell^-1^) | 35.3 |
| DD+DT (pg cell^-1^) | 5.1 |
| β-car (pg cell^-1^) | 2.1 |
| C (pg µm^3^) | 0.57 |
| C : N | 2.45 |
| Chl *a* : C | 17.5 |
| Daily C production (ng cell^-1^ day^-1^) | -0.13 |
| rETRm | 38 |
| Ek (µmol photons m^-2^s^-1^) | 68 |
| Eopt (µmol photons m^-2^s^-1^) | 140 |
| Y(NPQm) | 0.60 |
| Y(E50_NPQ_) (µmol photons m^-2^s^-1^) | 225 |
| Y(E50_NPQ_)/Eopt | 1.48 |

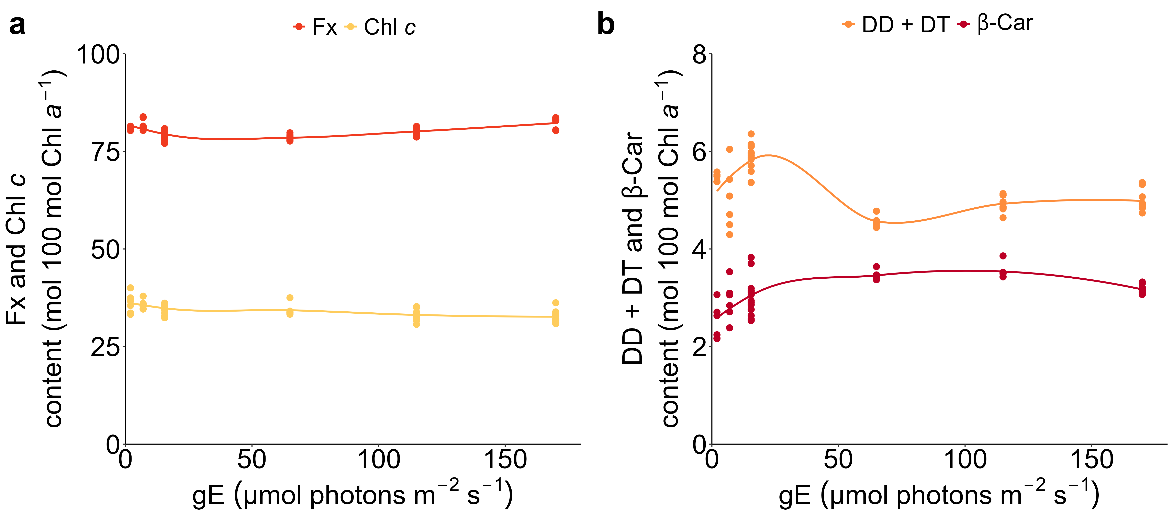

**Figure S7**: Pigment contents in mol 100 mol Chl a^-1^ of *P. strigosum* grown under increasing green-blue light intensities (gE). Pigments are (**a**) fucoxanthin (Fx), chlorophyll *c* (Chl *c*), and (**b**) the xanthophyll cycle pigments (DD+DT, diadinoxanthin and diatoxanthin), β-carotene (β-Car). Dots are raw data. Detailed statistical results for each pigment are shown in **Table S6**.

**Table S6**: Results from one-way ANOVA or Kruskal-Wallis and *post hoc* (pairwise Student’s t-test or Wilcoxson rank sum test) tests of pigment ratios in mol 100 mol Chl *a*^-1^ of *P. strigosum* grown under increasing green-blue light intensities (gE). Differences were considered significant for *p-value* < 0.050 (bold values).

| Fx | Kruskal-Wallis test (*p value* < 0.001) and pairwise Wilcoxon test | | | | |  |
| --- | --- | --- | --- | --- | --- | --- |
|  | gE | 2 | 7 | 15 | 62 | 115 |
|  | 7 | 0.5720 |  |  |  |  |
|  | 15 | **2.96E-02** | **1.90E-04** |  |  |  |
|  | 62 | **5.95E-03** | **6.80E-05** | 0.5720 |  |  |
|  | 115 | 0.5720 | **3.76E-02** | 0.3180 | 0.0668 |  |
|  | 170 | 0.1849 | 0.5720 | **2.20E-06** | **2.00E-06** | **2.67E-03** |
| Chl *c* | ANOVA test (*p value* < 0.001) and pairwise student test | | | | |  |
|  | gE | 2 | 7 | 15 (GBL) | 62 (GBH) | 115 |
|  | 7 | 1.0000 |  |  |  |  |
|  | 15 | 0.7510 | 1.0000 |  |  |  |
|  | 62 | 0.5107 | 1.0000 | 1.0000 |  |  |
|  | 115 | **1.28E-02** | **4.35E-02** | 0.1948 | 0.8539 |  |
|  | 170 | **2.90E-03** | **1.16E-02** | **4.35E-02** | 0.5107 | 1.0000 |
| DD+DT | Kruskal-Wallis test (*p value* < 0.001) and pairwise Wilcoxon test | | | | |  |
|  | gE | 2 | 7 | 15 (GBL) | 62 (GBH) | 115 |
|  | 7 | 0.8983 |  |  |  |  |
|  | 15 | **1.73E-02** | 0.0553 |  |  |  |
|  | 62 | **1.73E-02** | 1.0000 | **9.60E-04** |  |  |
|  | 115 | **4.80E-03** | 1.0000 | **6.00E-05** | **7.99E-03** |  |
|  | 170 | **4.80E-03** | 1.0000 | **6.00E-05** | 0.0080 | 1.0000 |
| β-Car | Kruskal-Wallis test (*p value* < 0.001) and pairwise Wilcoxon test | | | | |  |
|  | gE | 2 | 7 | 15 (GBL) | 62 (GBH) | 115 |
|  | 7 | 0.2597 |  |  |  |  |
|  | 15 | 0.1757 | 0.6388 |  |  |  |
|  | 62 | **2.38E-02** | 0.2468 | 0.1506 |  |  |
|  | 115 | **1.40E-02** | 0.2448 | 0.0635 | 0.2751 |  |
|  | 170 | **5.60E-03** | 0.2478 | 0.2597 | **5.60E-03** | **2.60E-03** |

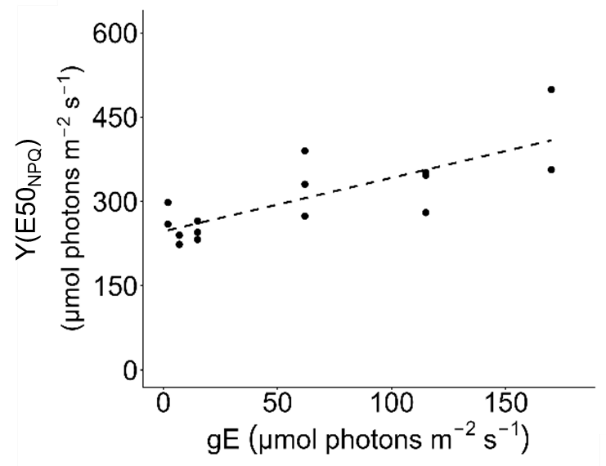

**Figure S8**: Y(E50_NPQ_), the light intensity for which the quantum yield of the regulated non-photochemical quenching (Y(NPQ)) reaches half-maximal expression *versus* growth light intensity (gE) for *P. strigosum* grown under increasing green-blue light intensities. Dots are raw data; line is a linear regression (R² = 0.61). Pearson correlation test was performed, *p-value* < 0.001, r = 0.78.

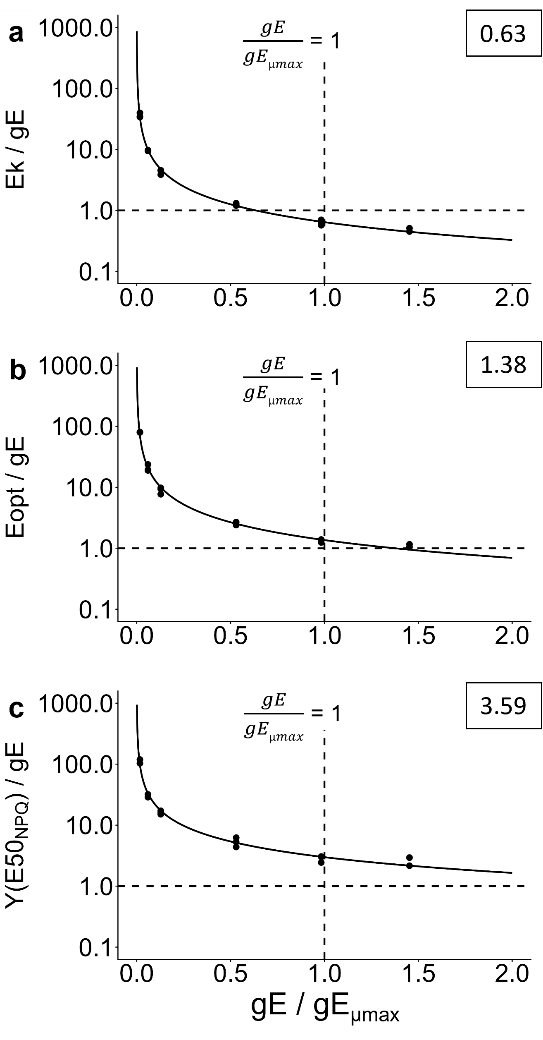

**Figure S9**: Ratios of light use optima (see **Table S1** for definitions) and the growth light intensity (gE), for *P. strigosum* grown under increasing green-blue light intensities. Ratios are shown as a function of the ratio gE/gE, with gE_µmax_ = the gE for the maximal growth rate. Dots are raw data, solid lines are power regression models fitted to the raw data R² are (**a**) 1, (**b**) 0.99 and (**c**) 0.98. Horizontal and vertical dashed lines indicate 1:1 ratio. The squared value on the top right represents the gE/gE_µmax_ at which the ratio of light use parameters crosses the horizontal 1:1 ratio.

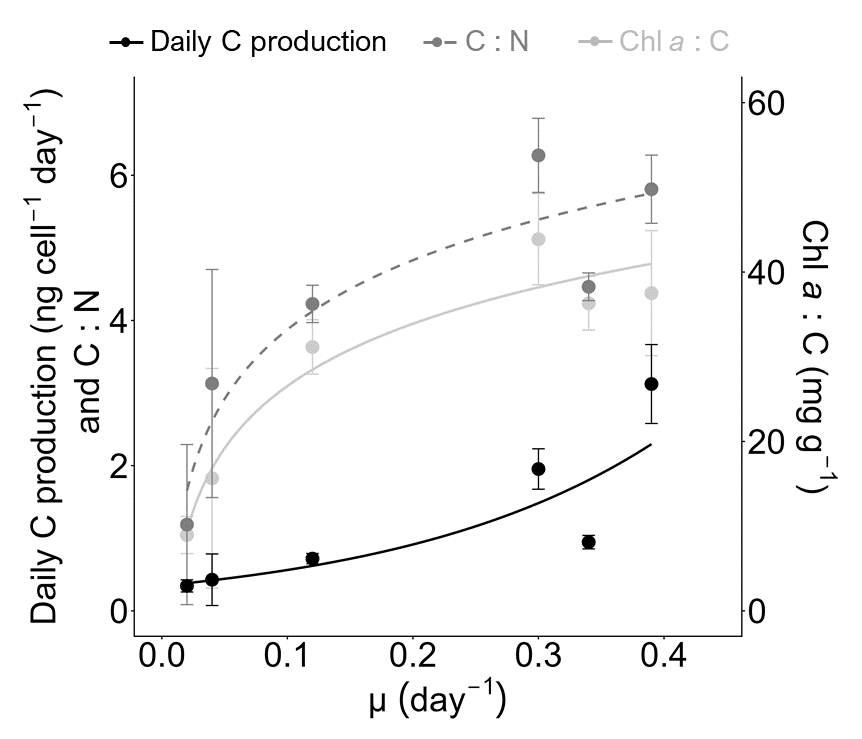

**Figure S10**: Daily C production (in ng cell^-1^day^-1^), Chl *a* : C (in mg g^-1^) and C : N versus growth rate for *P. strigosum* grown under increasing green-blue light intensities. Data are expressed as mean ± standard deviation. Lines are exponential (for daily C production, R² = 0.83) and logarithmic (for Chl *a* : C, R² = 0.93 and C : N, R² = 0.83) regressions.
